# A rapid, genetically-tractable human iPSC-derived model for studying tissue-resident macrophage biology in Alzheimer’s and other diseases

**DOI:** 10.64898/2026.10.08.757653

**Authors:** Hyo Lee, Grace B. Peppler, Michael Sewell, Alexandra Münch, Anna Podleśny-Drabiniok, Jeanne J. Kim, Joshua Orrick, Edoardo Marcora, Alison M. Goate

## Abstract

Large-scale genetic and preclinical studies strongly implicate myeloid cells, particularly brain-resident macrophages, in the pathogenesis of Alzheimer’s disease (AD). Efferocytic clearance of apoptotic cells and other cholesterol-rich cellular debris is a core macrophage function and represents an active area of AD research. However, human tissue-resident macrophages remain difficult to model *in vitro* as they acquire their transcriptional and functional identities in response to signals from their host tissue microenvironment. Many existing differentiation protocols mimic human brain macrophage developmental trajectories, but these protocols rely either on multi-step cytokine cocktails and lengthy differentiation or transduction of multiple transcription factors, both of which introduce variability and complicate genetic manipulation. To address these limitations, we developed a simplified protocol that generates high-purity (>90% CD45+CD11b+) human iPSC-derived macrophage-like cells (iMacs) from cryopreservable erythro-myeloid progenitor (EMP)-like intermediates in 19 days, including a final 7-day maturation step driven by macrophage colony-stimulating factor (M-CSF) alone. Transcriptomic profiling reveals that iMacs show enrichment for phagocytic and lysosomal programs, processes integral to efferocytosis. At both the transcriptional and protein level, iMacs have higher expression of scavenger receptors such as *CD36*, *CD163* and *MRC1*, and of AD risk genes such as *TREM2* and *APOE*. Functionally, iMacs show enhanced phagocytic uptake of diverse substrates relative to iPSC-derived microglia, including carboxylated latex beads, zymosan, amyloid-β 1-42, and amyloid-β 1-40, and exhibit expanded lysosomal mass. Robust TREM2 expression and signaling make iMacs a sensitive platform for interrogating TREM2 biology, with markedly higher phosphorylation of SYK and PLCγ2 in response to TREM2 engagement compared to iPSC-derived microglia. iMacs also tolerate lipid-mediated siRNA transfection with high survival and knockdown efficiency, enabling rapid genetic perturbation of AD risk genes. Together, these features make iMacs a practical, scalable, and genetically tractable human macrophage model for studying the regulation and cell biology of core macrophage functions that are known to play critical roles in AD, aging, and other disorders.

## Introduction

Central nervous system (CNS)-resident macrophages are increasingly recognised as active contributors to neurodegenerative diseases(Van Hove et al. 2025). Their contribution is particularly evident in Alzheimer’s disease (AD), where human genetics strongly implicates macrophages in disease risk. Genome-wide association studies (GWAS) have identified many common AD risk variants that are enriched in enhancers active in myeloid cells. Among those myeloid cells are microglia, the primary resident macrophages of the brain (Huang et al. 2017; Nott et al. 2019; Novikova et al. 2021). Additionally, many nominated risk genes, including *TREM2*, *PLCG2*, *INPP5D* and *MS4A4A*, are expressed predominantly in macrophages (Jansen et al. 2019; Bellenguez et al. 2022). Pathway analyses of AD GWAS further implicate processes critical to the core macrophage function of efferocytosis, including cholesterol metabolism, endocytosis and innate immunity (Askarova et al. 2025; Jones et al. 2010; Kunkle et al. 2019). Efferocytosis is the mechanism by which phagocytes recognize, engulf, and degrade apoptotic cells and other cholesterol-rich cellular debris, which represents a critical component of their metabolic adaptation to either store or eliminate digested material. This process is essential for maintaining immune tolerance, promoting tissue repair, and preserving tissue homeostasis throughout the lifespan (Romero-Molina et al. 2022).

Single-cell studies have identified microglial transcriptomic states closely associated with efferocytosis, most notably disease-associated microglia (DAM). DAM are known to appear in neurodegenerative conditions that include AD and aging. This cell state arises in response to the hallmarks of these degenerative conditions, which include dystrophic neurites surrounding amyloid-β plaques, apoptotic cells, synaptosomes, and myelin debris. DAM cluster around amyloid plaques and upregulate genes involved in phagocytosis, lysosomal degradation, and cholesterol efflux genes, including the AD risk genes *APOE* and *TREM2* (Keren-Shaul et al. 2017; Krasemann et al. 2017; De Schepper et al. 2023; Lan et al. 2024; Barclay et al. 2024). TREM2 signaling through SYK is critical for activating this efferocytic program, as TREM2- and SYK- deficient microglia fail to clear myelin debris and have impaired clustering around amyloid plaques. In addition, microglia with the AD risk variant TREM2 R47H do not fully activate the DAM response (Poliani et al. 2015; Cignarella et al. 2020; Nugent et al. 2020; Y. Wang et al. 2015; S. Wang et al. 2022). Recent literature has implicated the historically understudied border-associated macrophages (BAMs) in AD pathogenesis. These cells reside in the perivascular spaces, meninges and choroid plexus, and largely arise from the same embryonic yolk-sac erythro-myeloid progenitors (EMPs) as microglia (Goldmann et al. 2016; Masuda et al. 2022; Utz et al. 2020; Brioschi et al. 2023). BAMs are also highly phagocytic, expressing high levels of scavenger receptors such as *CD163*, *CD36* and *CD206*, as well as AD risk genes such as *APOE* (Van Hove et al. 2019; Sankowski et al. 2024). Loss of BAMs has been found to worsen amyloid plaque burden in mouse models (Drieu et al. 2022; Adler et al. 2026).

Much of our understanding of these macrophage states derives from animal models. However, microglial gene expression patterns diverge across species, particularly for those implicated in neurodegenerative disease. Human DAM shows only partial overlap with DAM in mice (Gosselin et al. 2017; Geirsdottir et al. 2019; Gerrits et al. 2021; Sun et al. 2023; Mancuso et al. 2024), which highlights the importance of using human systems for studying disease-relevant macrophage biology. *In vitro* systems are particularly suited to the study of efferocytosis, as they enable functional assays that assess substrate uptake and lysosomal degradation that can be challenging to measure *in vivo*. Moreover, *in vitro* systems are more susceptible to direct genetic perturbation than animal models, which marks them as an indispensable tool in the study of AD risk genes (Maguire et al. 2022). However, primary human microglia are difficult to obtain and rapidly lose their identity in culture (Gosselin et al. 2017). Consequently, induced pluripotent stem cell (iPSC)-derived myeloid cells have emerged as an attractive alternative for modeling human CNS macrophages *in vitro*.

Recapitulating disease-associated states in these models continues to represent a challenge in the field. DAM-like profiles have been induced in iPSC-derived microglia (iMGLs) through exposure to efferocytic substrates like apoptotic neurons and synapses, or by engineering cell lines to overexpress or delete transcription factors that regulate this state (Dolan et al. 2023; Podleśny-Drabiniok et al. 2024; Podlesny-Drabiniok et al. 2026). BAM-like cells have likewise been generated by omitting TGFβ1, a cytokine indispensable for microglial but not for BAM development (Butovsky et al. 2014; Utz et al. 2020; Brioschi et al. 2023) from an iMGL protocol (Barr et al. 2025). However, this BAM protocol requires long culture schedules and multiple costly cytokine supplements, maintaining major obstacles of traditional iMGL protocols (Vasoya et al. 2026; Haenseler et al. 2017; Washer et al. 2022; Abud et al. 2017; McQuade et al. 2018). These lengthy protocols can increase batch-to-batch variability, raise spending, and limit throughput, while the resulting cells can be difficult to manipulate genetically. Though faster approaches have been developed, these rely on transcription factor transduction and engineered, inducible lines (Dräger et al. 2022; Liu et al. 2025). We have previously established a macrophage model of efferocytosis using the THP1 cell line that is more accessible and readily manipulated than iMGLs (Podleśny-Drabiniok et al. 2025), but these cells are of monocytic lineage and therefore do not recapitulate the embryonic origin of the majority of CNS and other tissue-resident macrophages.

Here we describe a simplified 19-day protocol that generates human iPSC-derived macrophages (iMacs) from cryopreservable EMP-like progenitors, using a final 7-day maturation step driven by M-CSF alone. iMacs share transcriptional features with specific CNS macrophage states, including BAMs and DAM, and have elevated expression of genes involved in efferocytosis such as the surface receptors *CD36, CLEC7A, CD163* and *CD206*. Additionally, iMacs exhibit elevated expression of key AD risk genes such as *TREM2* and of genes implicated in lysosomal function. Functionally, iMacs show enhanced uptake of diverse substrates, increased lysosomal content, and a markedly enhanced signaling in response to TREM2 stimulation relative to iMGLs. iMacs also tolerate siRNA-mediated knockdown across multiple donor lines without significant loss of viability, enabling rapid perturbation of the genes regulating these processes. Together, these features make iMacs a practical, scalable, and genetically tractable human tissue-resident macrophage model for studying AD-relevant pathways such as efferocytosis, providing a vital tool to support functional genomics research and target discovery for Alzheimer’s and other diseases.

## Results

### Single-Cytokine Maturation of HPCs Efficiently Differentiates iPSCs into iMacs

To generate iMacs, human iPSCs were differentiated into erythro-myeloid progenitor (EMP)-like haematopoietic progenitor cells (HPCs) over 12 days using commercial media (STEMdiff Media A and B) as described previously (Abud et al. 2017; McQuade et al. 2018). These precursors were subsequently differentiated into iMacs over a further 7 days using commercially available macrophage media supplemented with M-CSF alone. In contrast, iMGL protocols supplement the same HPC stage with factors that mimic the brain microenvironment, including TGF-β and IL-34 to establish homeostatic microglial identity (Y. Wang et al. 2012; Butovsky et al. 2014; Utz et al. 2020) and the neuronal/astrocytic-derived ligands CX3CL1 and CD200 over a considerably longer maturation period (Abud et al. 2017; McQuade et al. 2018) (**Fig. 1a**). We reasoned that omitting these microglial identity and maturation signals, in particular TGF-β and IL-34, whose loss reprograms microglia towards border-associated and disease-associated states (Kana et al. 2019; Bedolla et al. 2024; McKinsey et al. 2025; Devlin et al. 2025; Hernandez-Rasco et al. 2026; Krasemann et al. 2017; Utz et al. 2020; Brioschi et al. 2023), particularly those characterized by increased expression of genes with key roles in efferocytosis (e.g., DAM and lipid-associated macrophage (LAM) states). iMacs exhibited round, amoeboid morphology under brightfield imaging, whereas iMGLs were more elongated and ramified (**Fig. 1b**). The resulting cultures were highly pure (>90% CD45⁺CD11b⁺) by day 19, with viability and reproducibility comparable to iMGLs across multiple genetic backgrounds (**Fig. 1c**; **Supp. Fig. 1a–b**).

**Figure 1.**
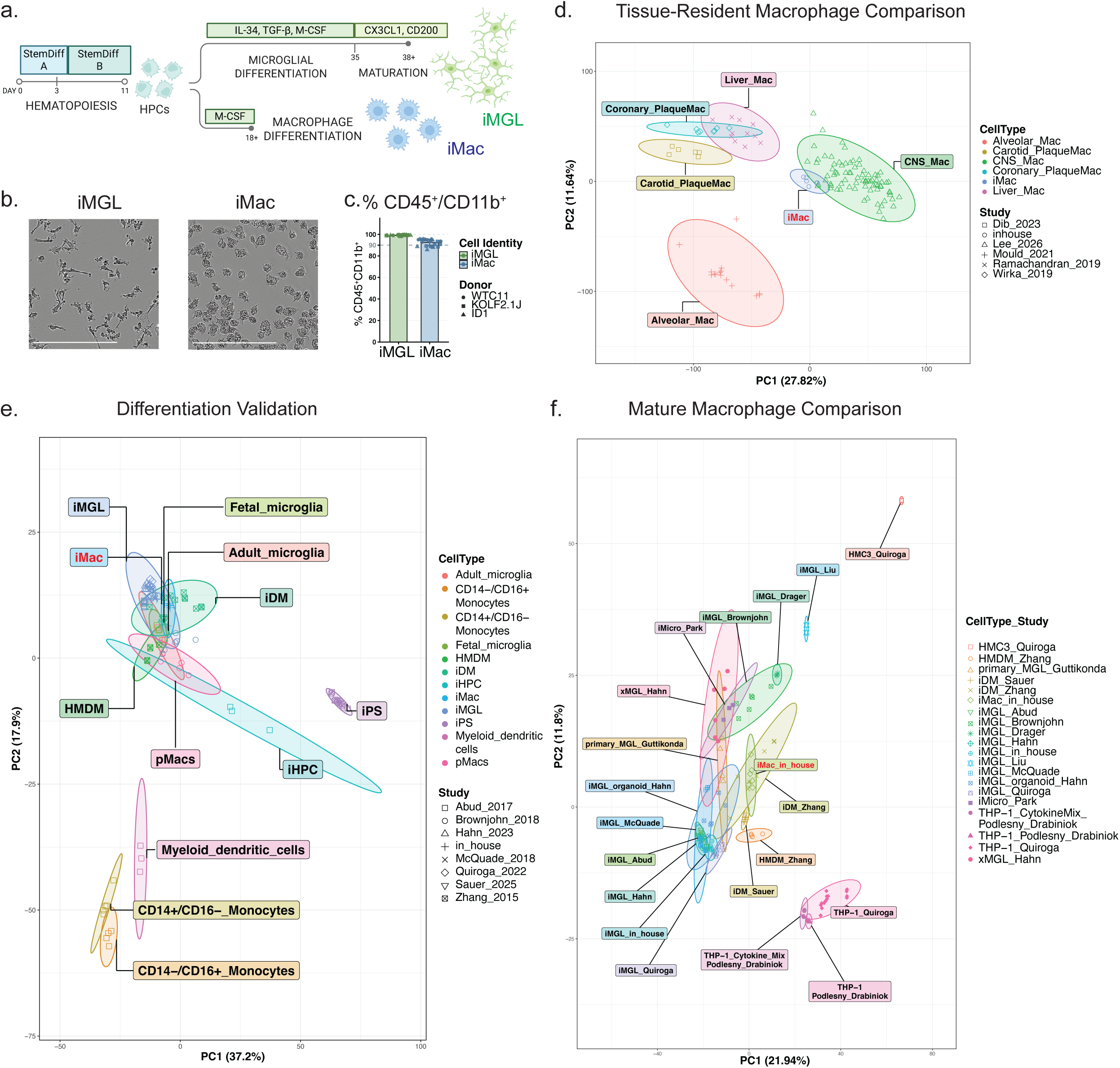
Rapid generation of human iPSC-derived macrophages (iMacs). (a) Schematic of the differentiation protocols showing shared initial generation of hematopoietic progenitor cells (HPCs) from iPSCs, followed by differentiation into iMGLs (28 days) or iMacs (7 days). (b) Representative brightfield images of iMGLs and iMacs. Scale bar = 200um. (c) Quantification of myeloid cell purity (% CD45⁺ CD11b⁺ cells) in iMGLs and iMacs across 3 differentiations, 3 iPSC donor lines (WTC11, KOLF2.1J, and ID1).(d) Principal component analysis (PCA) of RNA-seq data comparing iMacs against primary human tissue-resident macrophages from other published studies (Methods). Ellipses indicate the region expected to contain 95% of samples in each group, assuming a bivariate normal distribution. (e) Principal component analysis (PCA) of RNA-seq data comparing iMacs against iPSCs, iPSC-derived hematopoietic progenitor cells (iHPCs) and other cell types including primary adult and fetal human microglia, and dendritic cells from other published studies (Methods). **(f)** PCA comparison across mature *in vitro* macrophage and microglial models, including in-house iMacs and iMGLs ((WTC11 donor line, n = 3 clones, 2 differentiations per cell type), and primary human microglia. Ellipses indicate the region expected to contain 95% of samples in each group, assuming a bivariate normal distribution.

Tissue-resident macrophages (TRMs) arise from common EMP-derived precursors and diverge into tissue-specific transcriptional identities, shaped by their local microenvironment (Gosselin et al. 2014; Mass et al. 2016). Given that iMacs mature without the brain-derived cues used in iMGL protocols, we tested whether they more closely resemble CNS or peripheral tissue-resident macrophage populations. Bulk RNA sequencing was performed on iMacs differentiated from three independent WTC11 iPSC clones, and the resulting profiles were benchmarked against primary human TRMs from diverse organs, identified through re-analysis of published scRNA-seq datasets. These included macrophages from liver (Ramachandran et al. 2019), lung (Mould et al. 2021), and CNS (both microglia and BAM) (Lee et al. 2026) of control subjects, as well as macrophages from carotid (Dib et al. 2023) and coronary artery (Wirka et al. 2019) atherosclerotic plaques. We compared overall transcriptomic distance by PCA visualization (**Fig. 1d**) and quantification of Euclidean distance and Kullback–Leibler divergence metrics (Bobadilla-Suarez et al. 2020) (**Supp. Fig. 2a–b**). iMacs were closest to CNS macrophages and furthest from alveolar macrophages, which showed higher expression of established identity genes such as *MARCO*, *PPARG* and *FABP4* (Bain and MacDonald 2022) (**Supp. Fig. 2c**). Liver and atherosclerotic plaque macrophages clustered together, which may reflect shared lipid-handling LAM programs arising from their lipid-rich tissue environments (Jaitin et al. 2019; K. Kim et al. 2022; Dib et al. 2023). iMacs also expressed lower levels of hepatic and coronary macrophage markers such as *TIMD4* and *MMP13* (W. Zhang et al. 2019; Nusse and Kubes 2025), and, like iMGLs, higher levels of CNS macrophage markers shared by microglia and BAM such as *CX3CR1* and CD11b (*ITGAM*) (Dermitzakis et al. 2023).

### iMacs cluster with primary and stem cell derived human macrophages and microglia rather than progenitor or monocyte populations

Given their closer transcriptomic similarity to CNS macrophages, we next asked how closely iMacs resemble microglia. We performed a PCA comparing iMacs with published datasets of primary and *in vitro* microglial and macrophage models that included other myeloid cells such as monocytes and dendritic cells, as well as intermediate stages of the *in vitro* differentiation protocols (Abud et al. 2017; Brownjohn et al. 2018; McQuade et al. 2018; Quiroga et al. 2022; Sauer et al. 2025; H. Zhang et al. 2015; Han et al. 2023) (**Fig. 1e**). iMacs were clearly distinct from circulating monocytes, dendritic cells, iPSCs, and iPSC-derived haematopoietic progenitors, with primitive macrophages (pMac) lying at the edge of the mature cluster. Instead, iMacs clustered with fetal and adult primary microglia, alongside other *in vitro* macrophage models (HMDMs, iDMs and iMGLs).

This apparent similarity to microglia was unexpected given that iMacs are matured without the brain-derived cues used in iMGL protocols. However, since the first comparison included iPSCs, progenitors, monocytes, and dendritic cells, its principal components were largely driven by differentiation stage and by the separation of tissue-resident from circulating myeloid cells, leaving little resolution among the mature models themselves. To better discriminate between these models, we performed a second PCA comparing iMacs solely with mature microglial and macrophage models, including iMGLs generated by transcription factor-driven (iTF) protocols(Dräger et al. 2022; Liu et al. 2025), microglia matured within cerebral organoids or after xenotransplantation(Park et al. 2023; Han et al. 2023), cultured primary microglia(Guttikonda et al. 2021), the immortalised HMC3 and THP-1 lines(Quiroga et al. 2022; Podleśny-Drabiniok et al. 2025), and the iDMs, HMDMs and iMGLs from **Fig. 1e**. For each model, we then quantified its Euclidean distance and Kullback–Leibler (KL) divergence from iMacs (**Fig. 1f; Supp. Fig. 3a–b**). By both metrics, iMacs were most similar to other iPSC-derived (iDMs) and monocyte-derived macrophages (HMDMs) (Sauer et al. 2025; H. Zhang et al. 2015), and most divergent from the immortalised THP-1 and HMC3 lines and from iTF-driven iMGLs. Notably, iMacs showed closer similarity to macrophage models than to any of the microglial models, including iMGLs generated in parallel in our lab. This indicates that omitting the microglial maturation cues yields a macrophage state transcriptionally distinct from homeostatic microglia.

### iMacs overlap transcriptionally with disease-associated macrophage states and show elevated expression of BAM scavenger receptors and certain AD risk genes

Having established that iMacs are transcriptionally distinct from iMGLs, we performed differential expression analysis between the two to identify gene expression changes underlying this difference. Differential expression analysis showed that relative to iMGLs, iMacs display upregulation of scavenger receptors which are enriched in other tissue resident macrophages such as BAM, including the scavenger receptors *F13A1* (logFC = 12.6, *P*_adj_ = 4.4 × 10^−6^), *MRC1* (logFC = 4.8, *P*_adj_ = 1.5 × 10^−5^), *CD36* (logFC = 5.0, *P*_adj_ = 1.2 × 10^−6^), *CD163* (logFC = 2.1, *P*_adj_ = 8.0 × 10^−4^) and *PF4* (logFC = 1.8, *P*_adj_ = 0.019), alongside lower expression of homeostatic microglial genes such as *CX3CR1* (logFC = −5.2, *P*_adj_ = 1.6 × 10^−6^), *NAV3* (logFC = −4.4, *P*_adj_ = 4.0 × 10^−4^), *P2RY12* (logFC = −2.7, *P*_adj_ = 8.5 × 10^−4^) and *CST3* (logFC = −2.3, *P*_adj_ = 2.9 × 10^−6^) (**Fig. 2a**; **Supplementary Table 1**) (Sankowski et al. 2024; Mass et al. 2016). iMacs also showed marked downregulation of MHC class II antigen-presentation genes, including *HLA-DRA* (logFC = −9.5, *P*_adj_ = 2.2 × 10^−6^), *HLA-DRB1* (logFC = −8.2, *P*_adj_ = 2.1 × 10^−4^), *HLA-DPA1* (logFC = −6.8, *P*_adj_ = 4.8 × 10^−7^), *HLA-DPB1* (logFC = −5.8, *P*_adj_ = 6.5 × 10^−5^), *HLA-DRB5* (logFC = −5.6, *P*_adj_ = 2.1 × 10^−5^), *HLA-DQB1* (logFC = −2.5, *P*_adj_ = 0.013), *HLA-DMA* (logFC = −2.1, *P*_adj_ = 5.1 × 10^−5^) and *HLA-DMB* (logFC = −1.2, *P*_adj_ = 6.8 × 10^−5^), which are characteristic markers of specific BAM subpopulations, notably those of the dura mater (Vara-Pérez and Movahedi 2025; Van Hove et al. 2019). No difference in expression was also observed for the canonical perivascular marker LYVE1 (logFC = −0.5, FDR = 0.42) with low expression across samples (mean log2 CPM = 0.33), indicating that while iMacs share features with BAMs, they do not fully recapitulate profiles of any single BAM subpopulation. Flow cytometry confirmed that several of our differential expression observations were recapitulated at the protein level, with increased surface expression of the scavenger receptors CD206 (encoded by *MRC1*; β = 14,454 gMFI, 95% CI: 8,877–22,984, *P*_adj_ = 0.002) and CD163 (β = 22,512 gMFI, 95% CI: 18,201–24,872, *P*_adj_ = 1.3 × 10^−4^), and reduced expression of the MHC class II marker HLA-DR (β = −10,988 gMFI, 95% CI: −15,277 to −4,403, *P*_adj_ = 0.021), relative to iMGLs (**Fig. 2b**).

**Figure 2.**
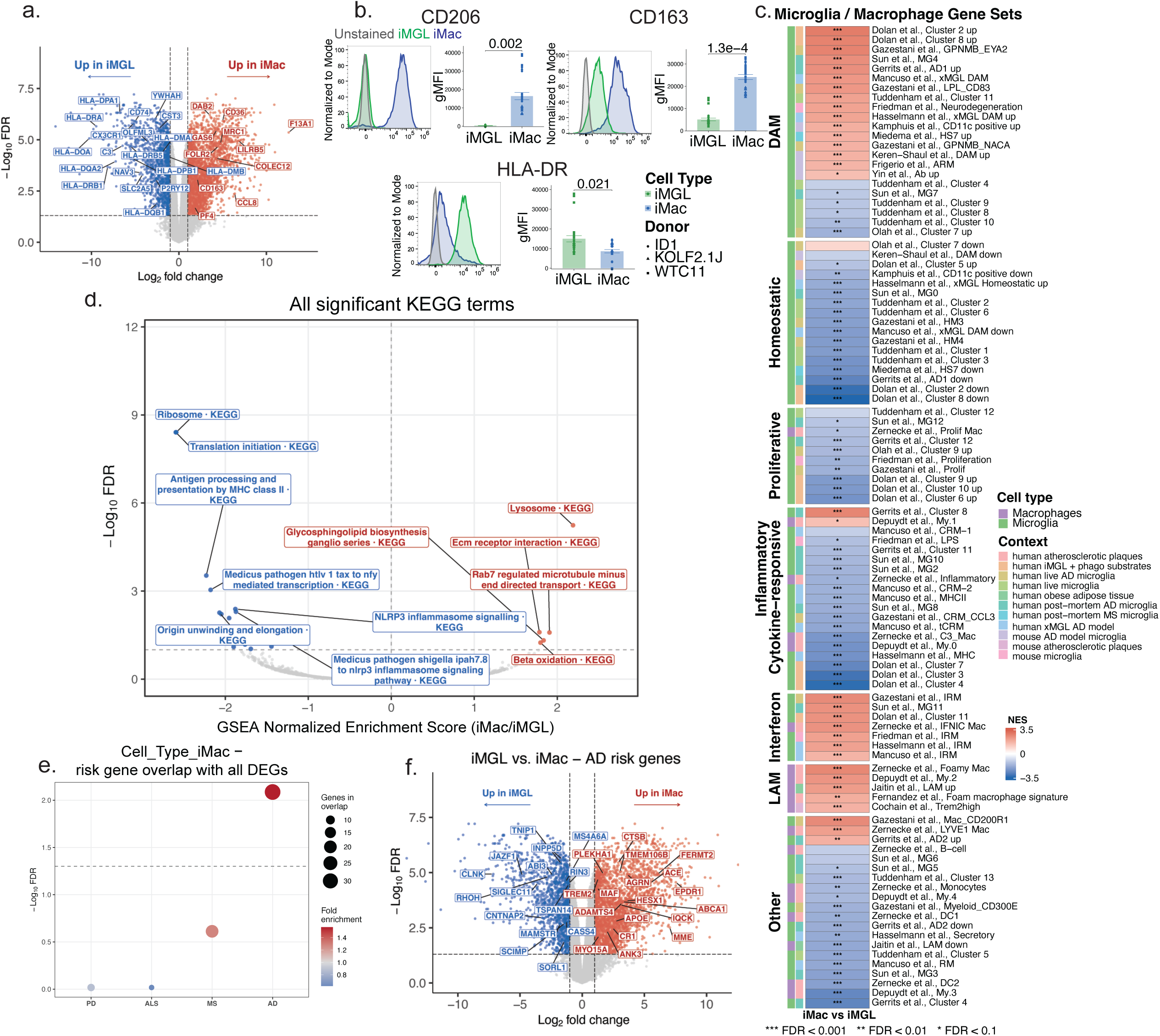
iMacs show strong enrichment for disease-associated microglia (DAM) pathway signatures and surface proteins. **(a)** Volcano plot of differentially expressed genes between iMacs and iMGLs (WTC11 donor line, n = 3 clones, 2 differentiations per cell type). Genes with FDR < 0.05 and |log_2_FC| > 1 are colored red (higher in iMacs) or blue (higher in iMGLs). **(b)** Representative flow cytometry histograms and geometric mean fluorescence intensity (gMFI) for CD206, CD163 and HLA-DR in iMGLs (green) and iMacs (blue), with unstained controls (gray). Data are mean ± SEM (n = 3 donor lines, 3 differentiations per donor, each point represents a well, with point shape indicating donor line (ID1, KOLF, WTC11). *P* values were obtained from linear mixed-effects models with donor and differentiation as fixed effects and a random intercept grouping replicate wells from the same donor line, cell type and differentiation, Holm-adjusted across all markers of the corresponding flow cytometry antibody panel. **(c)** Gene set enrichment analysis of published myeloid gene signatures against the iMac versus iMGL profile. Heatmap shows normalized enrichment scores (NES), grouped by state and annotated by the cell type and study context from which each signature was derived. **(d)** Volcano plot of KEGG pathway enrichment, plotted by NES and −log_10_ FDR. Colored points reached FDR < 0.1 (red, enriched in iMacs; blue, enriched in iMGLs). **(e)** Hypergeometric over-representation of risk genes for AD, PD, ALS and MS among all genes differentially expressed between iMacs and iMGLs. Point size shows the number of overlapping genes and color shows fold enrichment; the dashed line marks FDR = 0.05. **(f)** Volcano plot as in (a), with AD risk genes labeled. Red denotes AD risk genes higher in iMacs, blue denotes those higher in iMGLs. FDR, Benjamini–Hochberg-adjusted *P* value. Full details for statistical analyses of each panel are provided in the Supplementary Statistical Reporting Document

Unbiased gene set enrichment analyses (GSEA) against pathway databases revealed that iMGLs were enriched for pathways associated with translation (KEGG RIBOSOME, NES = −2.59, *P*_adj_ = 3.9 × 10^−9^; KEGG MEDICUS REFERENCE TRANSLATION INITIATION, NES = −2.60, *P*_adj_ = 3.9 × 10^−9^), as well as immune phenotypes such as GOBP ADAPTIVE IMMUNE RESPONSE (NES = −1.88, *P*_adj_ = 3.6 × 10^−7^), GOBP LYMPHOCYTE ACTIVATION (NES = −1.79, *P*_adj_ = 2.7 × 10^−9^), KEGG MEDICUS REFERENCE NLRP3 INFLAMMASOME SIGNALING PATHWAY (NES = −1.88, *P*_adj_ = 0.0041), and KEGG MEDICUS REFERENCE ANTIGEN PROCESSING AND PRESENTATION BY MHC CLASS II MOLECULES (NES = −2.23, *P*_adj_ = 3.0 × 10^−4^) (**Fig. 2c**; **Supp. Fig. 4a–d; Supplementary Table 2**), in keeping with our differential expression analysis showing downregulation of MHC class II genes. In contrast, iMacs showed enrichment for processes related to metabolism (KEGG GLYCOSPHINGOLIPID BIOSYNTHESIS GANGLIO SERIES, NES = 1.84, *P*_adj_ = 0.048; GOBP CELLULAR CATABOLIC PROCESS, NES = 1.48, *P*_adj_ = 3.1 × 10^−4^), extracellular matrix regulation (KEGG ECM RECEPTOR INTERACTION, NES = 1.78, *P*_adj_ = 0.025; REACTOME EXTRACELLULAR MATRIX ORGANIZATION, NES = 1.73, *P*_adj_ = 0.0013), and endolysosomal function (KEGG LYSOSOME, NES = 2.19, *P*_adj_ = 5.7 × 10^−6^; KEGG MEDICUS REFERENCE RAB7 REGULATED MICROTUBULE MINUS END DIRECTED TRANSPORT, NES = 1.91, *P*_adj_ = 0.026; GOCC ENDOSOME, NES = 1.45, *P*_adj_ = 0.0012; GOCC ENDOSOME MEMBRANE, NES = 1.56, *P*_adj_ = 0.0012).

Endolysosomal pathways underpin core macrophage clearance functions, including those of DAM. We therefore reasoned that iMacs may show strong transcriptomic overlap with this disease-associated state. To investigate this further, we performed another GSEA against transcriptomic signatures of macrophage states of interest, mined from both mouse and human studies (**Fig. 2d; Supplementary Table 3**). iMacs showed positive enrichment for signatures of lipid-clearance-associated states, including DAM (16/22 gene sets) and peripheral LAM (5/5 gene sets), two states that we have previously shown to share common transcriptional regulators across species and tissues (Podleśny-Drabiniok et al. 2024). iMacs were also enriched for interferon-response signatures (7/7 gene sets) and perivascular macrophage signatures (Mac_CD200R1 Gazestani et al. 2023; LYVE1 Mac Zernecke et al. 2023) (Zernecke et al. 2023; Gazestani et al. 2023). Conversely, iMac were negatively enriched for the vast majority of microglial homeostatic (15/17 gene sets) and other states found in disease such as proliferative (9/10 gene sets) and inflammatory cytokine/response (16/19 gene sets), as well as those of other immune cells (Zernecke et al., B−cell, Zernecke et al., Monocytes, Gazestani et al., Myeloid_CD300E, Zernecke et al., DC1, Zernecke et al., DC2) (Zernecke et al. 2023; Gazestani et al. 2023), in keeping with our previous analyses in **Fig. 1e** .

Importantly, several DAM signature genes, as well as transcription factors thought to orchestrate the DAM response, are candidate AD risk genes (Romero-Molina et al. 2022; Keren-Shaul et al. 2017; Podleśny-Drabiniok et al. 2024; Podlesny-Drabiniok et al. 2026). Moreover, genetic and genomic studies have shown that AD risk variants, unlike those of other neurodegenerative and neurological disorders, are particularly enriched in enhancers active in microglia and other myeloid cells (Huang et al. 2017; Nott et al. 2019; Askarova et al. 2025). We therefore asked whether the iMac transcriptional program is enriched for AD risk genes nominated at risk loci relative to those for other neurodegenerative diseases. Hypergeometric over-representation analysis of all genes differentially expressed between iMacs and iMGLs (both up- and downregulated) showed selective enrichment for genes positionally mapped to AD risk loci (33 of 74 tested genes differentially expressed vs. 21.0 expected; 1.57-fold enrichment; OR = 2.04, 95% CI 1.25–3.32; FDR = 0.008) (Jansen et al. 2019; Kunkle et al. 2019; Bellenguez et al. 2022), whereas no significant overlap was observed for loci associated with Parkinson’s disease (PD) (J. J. Kim et al. 2024; Nalls et al. 2019), amyotrophic lateral sclerosis (ALS) (van Rheenen et al. 2021) or multiple sclerosis (MS) (International Multiple Sclerosis Genetics Consortium 2019) (**Fig. 2e; Supplemental Table 4**). The direction of change in AD risk genes also mirrored that seen in DAM, as iMacs showed increased expression of TREM2, APOE, and CTSB(**Fig. 2f)** (Martins-Ferreira et al. 2025; Deczkowska et al. 2018). Together these findings highlight the potential utility of our iMac model in studying the roles of genes and macrophage states implicated as causal drivers in AD pathogenesis.

### iMacs Demonstrate Enhanced Phagocytic Capacity and Expanded Lysosomal Mass

The enrichment of clearance-associated macrophage states in iMacs, together with elevated expression of receptors such as TREM2, CD163, and CD206 (**Fig. 2**), led us to hypothesize that iMacs may have enhanced clearance capacity relative to iMGLs. To explore this, we revisited our GSEA, focusing on clearance-associated pathways and the genes driving their enrichment (**Fig. 3a**; **Supp. Fig. 5a-b**; **Supplementary Table 2**). Beyond the endolysosomal terms described above, iMacs were enriched for lysosomal lumen and acidification terms (GOCC LYSOSOMAL LUMEN, NES = 2.09, *P*_adj_ = 2.3 × 10^−4^; GOBP LYSOSOMAL LUMEN ACIDIFICATION, NES = 2.02, *P*_adj_ = 0.0056) and for the response to amyloid-beta (GOBP CELLULAR RESPONSE TO AMYLOID BETA, NES = 1.95, *P*_adj_ = 0.018) (**Fig. 3a**). Lysosomal terms such as KEGG LYSOSOME were largely driven by genes encoding lysosomal hydrolases, including multiple cathepsins such as the AD risk gene *CTSB* and the DAM-associated *CTSD* (**Supp. Fig. 5a**).

**Figure 3.**
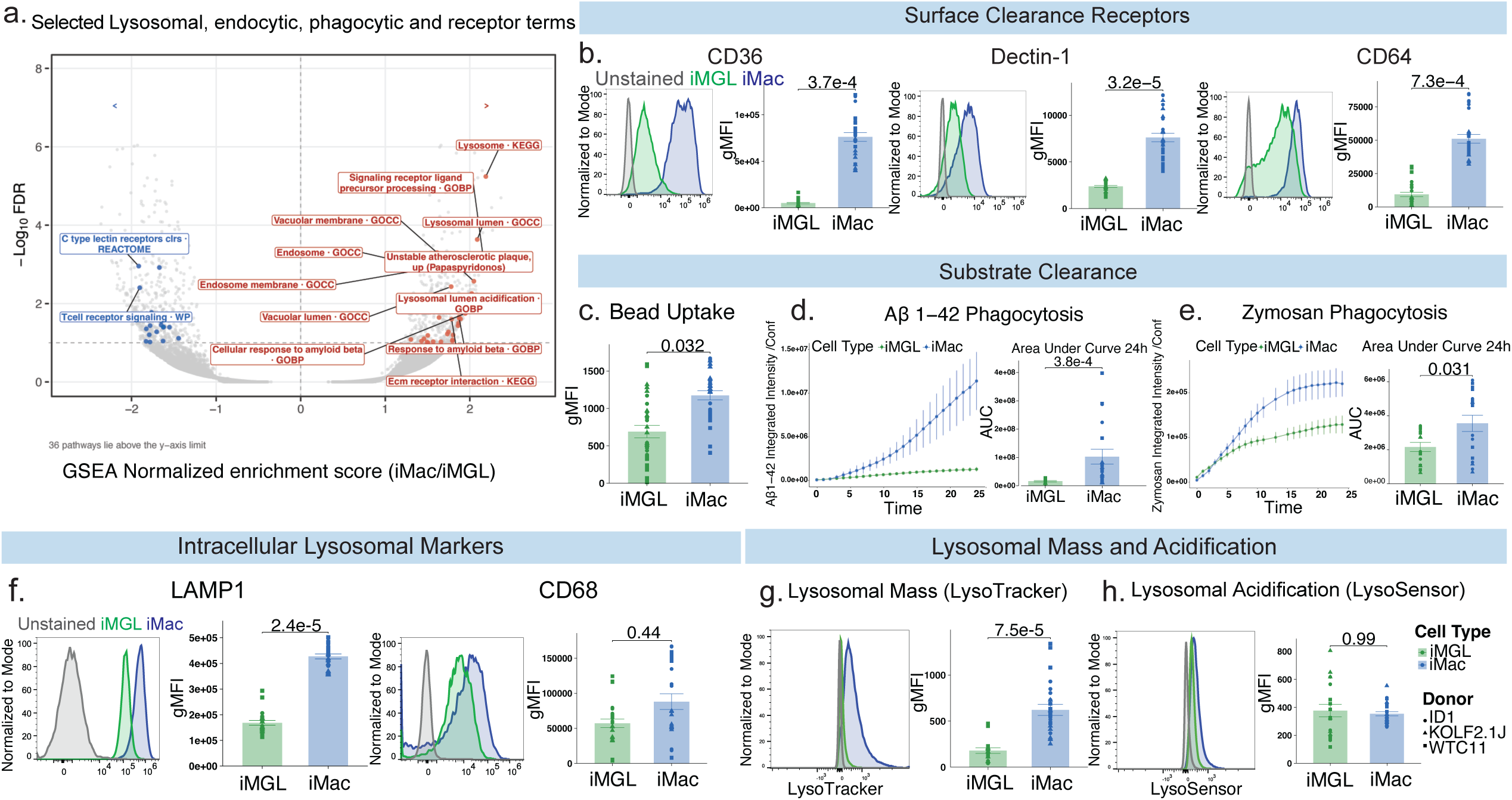
iMacs exhibit elevated scavenger receptor expression, enhanced phagocytic and lysosomal function compared to iMGLs. **(a)** Volcano plot of GSEA results showing normalized enrichment scores (iMac/iMGL) vs. -log_10_(FDR). Selected lysosomal, endocytic, phagocytic, and receptor-mediated pathways relevant to clearance function are highlighted. **(b)** Representative flow cytometry histograms and surface expression quantification (gMFI) for key scavenger receptors (CD36, CLEC7A, and CD64). **(c)** Fluorescent carboxylate latex beads uptake after 24 hours of treatment quantified by flow cytometry (gMFI). **(d)** Live-imaging phagocytosis time-course (left) and corresponding area under the curve (AUC) quantification (right) of 488-abeta42 total integrated intensity/confluence over 24 hours. **(e)** Live-imaging phagocytosis time-course (left) and AUC quantification (right) of pHrodo-labeled zymosan total integrated intensity/confluence over 24 hours. (f) representative flow cytometry histograms and intracellular expression quantification (gMFI) for lysosomal proteins LAMP1 and CD68, in iMGLs (green) and iMacs (blue) relative to unstained controls (gray). (g-h) Representative histograms and gMFI quantifications of lysosomal mass using LysoTracker Deep Red (left) and lysosomal pH/acidification using LysoSensor (right). Bars represent the mean and error bars of the SEM; each point represents one well, with point shape indicating donor line (ID1, KOLF, WTC11). Statistical significance was evaluated using linear mixed effects models. Full statistics are provided in the Supplementary Statistical Report.

In contrast, terms such as GOBP CELLULAR RESPONSE TO AMYLOID BETA were driven by genes encoding cell-surface receptors involved in substrate recognition, including *TREM2*, toll-like receptor 4 (*TLR4*), Fc gamma receptor IIb (*FCGR2B*) and *CD36* (Chen et al. 2025)(**Supp. Fig. 5b**). This suggests that iMacs may have an enhanced capacity to both sense and process substrates. Flow cytometry confirmed elevated surface expression of CD36 on iMacs relative to iMGLs (β = 77,495 gMFI, 95% CI: 62,159–85,403, *P*_adj_ = 3.7 × 10^−4^), and additionally revealed higher surface expression of the Fc gamma receptor CD64 (FcγRI; β = 40,247 gMFI, 95% CI: 33,811–53,052, *P*_adj_ = 7.3 × 10^−4^) (**Fig. 3b**). Interestingly, upregulation of substrate-recognition receptors was not universal across receptor families at the transcript level. Transcriptomic and pathway analyses indicated lower expression in iMacs of genes encoding C-type lectin receptors (CLRs), a family of pattern recognition receptors (PRRs), including *CLEC7A* (logFC = −1.4, *P*_adj_ = 8.1 × 10^−5^), *CLEC4A* (logFC = −2.7, *P*_adj_ = 0.0014) and *CLEC10A* (logFC = −5.2, *P*_adj_ = 9.8 × 10^−5^), shown in negative enrichment for REACTOME C TYPE LECTIN RECEPTORS CLRS (NES = −1.92, *P*_adj_ = 0.0011; **Fig. 3a**). *CLEC7A* (which encodes for Dectin-1) is also a canonical DAM marker (Krasemann et al. 2017; Barclay et al. 2024), which led us to ask whether this lower expression was mirrored at the protein level. Surprisingly, as with the other surface clearance receptors, Dectin-1 was significantly higher on iMacs than on iMGLs (β = 5,089 gMFI, 95% CI: 4,445–6,141, *P*_adj_ = 3.2 × 10^−5^; **Fig. 3b**). This may reflect post-transcriptional regulation across receptors, such as differences in protein half-life or receptor trafficking and recycling at the cell surface.

To determine whether these transcriptional and protein-level differences were accompanied by enhanced clearance capacity, we performed a suite of functional assays to assess overall phagocytic capacity, substrate-specific uptake, and lysosomal mass and acidification. To assess overall phagocytic capacity, cells were treated with carboxylated fluorescent latex beads, which are widely used as apoptotic-cell surrogates as their negative surface charge approximates phosphatidylserine exposure on dying cells. Cells were treated with beads for 24 hours and uptake was measured by flow cytometry, with fluorescence intensity peaks used to quantify bead number within each cell (**Supp. Fig. 6a-d**). iMacs had significantly enhanced phagocytosis of latex beads compared to iMGLs, with higher bead fluorescence (β = 346 gMFI, 95% CI: 56–635, *P*_adj_ = 0.032) and a higher proportion of bead-positive cells (39.2% vs 24.6%; OR = 1.98, 95% CI: 1.31–2.99, *P* = 0.001) (**Fig. 3c; Supplementary Fig. 6d**). This reflected a shift towards cells containing multiple beads, with iMacs showing more cells containing 2 (OR = 2.24, 95% CI: 1.45–3.46, *P*_adj_ = 0.001) or 3+ beads (OR = 2.89, 95% CI: 1.52–5.49, *P*_adj_ = 0.003), consistently across all three donor lines (**Supp. Fig. 6d**). Cells containing a single bead were also more frequent in iMacs (OR = 1.41, 95% CI: 1.03–1.93, *P*_adj_ = 0.031), although this difference was donor-dependent (cell type × donor interaction, *P* = 0.013) (**Supp. Fig. 6e**). Given the elevated surface expression of CD36 and Dectin-1 on iMacs (**Fig. 3b**), we next asked whether iMacs also showed enhanced uptake of substrates recognized by these receptors such as amyloid-beta (Aβ) and zymosan (Zhao et al. 2023; Silva et al. 2024; El Khoury et al. 2003; Doens et al. 2017). We measured uptake of pHrodo-labeled zymosan, AF488-conjugated Aβ 1-42, and AF488-conjugated Aβ 1-40 via live imaging over the course of 24 hours. Compared to iMGLs, iMacs had enhanced uptake of all tested substrates (**Fig. 3d–e; Supp Fig. 7**). Uptake of both Aβ species increased over time between cell types (cell type × time interaction: Aβ 1-42, χ²(3) = 102.9, *P* = 3.7 × 10^−22^; Aβ1–40, χ²(3) = 222.8, *P* = 4.9 × 10^−48^ ), with iMacs showing greater uptake from 8 h for Aβ 1-42 and from 16 h for Aβ 1-40 (**Fig 3e; Supp. Figure 7a-b; Supplementary Statistical Reporting Document**), and greater total uptake over 24 h measured by area under the curve (AUC) values (AUC fold change: Aβ1-42, 7.9, 95% CI: 4.4–14.3, *P*_adj_ = 3.8 × 10^−4^; Aβ1-40, 4.9, 95% CI: 2.5–9.5, *P*_adj_ = 0.042). Zymosan uptake similarly diverged from 8 h (cell type × time interaction, χ²(3) = 154.1, *P* = 3.4 × 10^−33^), with 1.6-fold greater total uptake in iMacs over the 24-hour period (95% CI: 1.1–2.5, *P* = 0.031) (**Fig 3e**).

Next, to determine whether the transcriptional enrichment of lysosomal genes in iMacs translated to increased protein expression, we evaluated the lysosomal markers LAMP1 and CD68. Both were significantly upregulated at the transcript level (*LAMP1*, logFC = 1.1, *P*_adj_ = 1.7 × 10^−4^; *CD68*, logFC = 1.9, *P*_adj_ = 2.4 × 10^−5^). While iMacs showed significantly elevated LAMP1 relative to iMGLs (β = 294,168 gMFI, 95% CI: 242,778–327,936, *P*_adj_ = 7.3 × 10^−4^; **Fig. 3f**), no significant difference was observed for CD68 (β = 16,693 gMFI, 95% CI: −8,865 to 76,074, *P*_adj_ = 0.44; **Fig. 3f**), which may partly reflect its greater variability between differentiations compared with LAMP1 (**Supplementary Statistical Report**).

We then performed functional assays assessing lysosomal mass and acidification, since lysosomal impairment is commonly implicated in many neurodegenerative disorders such as AD (De Tito and Tooze 2026). To measure lysosomal mass, we stained with LysoTracker Deep Red and assessed fluorescence using flow cytometry. iMacs had significantly higher lysosomal mass compared to iMGLs (3.9-fold; β = 587 gMFI, 95% CI: 369–871, *P*_adj_ = 7.5 × 10^−5^) (**Fig. 3g**). To assess lysosomal pH, we performed the same experiment using LysoSensor Green. There was no significant difference between cell types (fold change = 1.00, 95% CI: 0.74–1.36, *P*_adj_ = 0.998), although LysoSensor signal varied considerably between differentiations (**Fig. 3h; Supplementary Statistical Reporting Document**). Nonetheless, the increased lysosomal mass, higher expression of phagocytic surface receptors and lysosomal proteins, and enhanced uptake of all tested substrates together suggest that iMacs possess enhanced capacity for phagocytic uptake and lysosomal clearance.

### iMacs have increased TREM2 expression and enhanced TREM2 signaling compared to iMGLs

*TREM2* is another AD risk gene thought to play a critical role in efferocytosis(Romero-Molina et al. 2022) that has higher expression in iMacs than in iMGLs in our RNA-seq data (**Fig. 2f**). We therefore asked whether this was reflected in TREM2 protein levels and downstream signaling. Consistent with the enrichment of DAM gene signatures observed in our GSEA transcriptional profiling, *TREM2* was among the genes significantly upregulated at the RNA level in iMacs compared to iMGLs. At the protein level, western blot analysis of the cell membrane fraction demonstrated elevated expression of both full-length TREM2 (TREM2-FL; β = 11,513 a.u., 95% CI: 8,308–14,652, *P*_adj_ = 0.011) and the C-terminal fragment (TREM2-CTF; β = 10,857 a.u., 95% CI: 7,456–14,253, *P*_adj_ = 0.011) in iMacs compared to iMGLs (**Fig. 4a, b**), a finding supported by the robust and reproducible increase in surface TREM2 expression observed via flow cytometry (β = 12,347 gMFI, 95% CI: 8,207–13,846, *P*_adj_ = 7.8 × 10^−4^; **Fig. 4c**).

**Fig 4.**
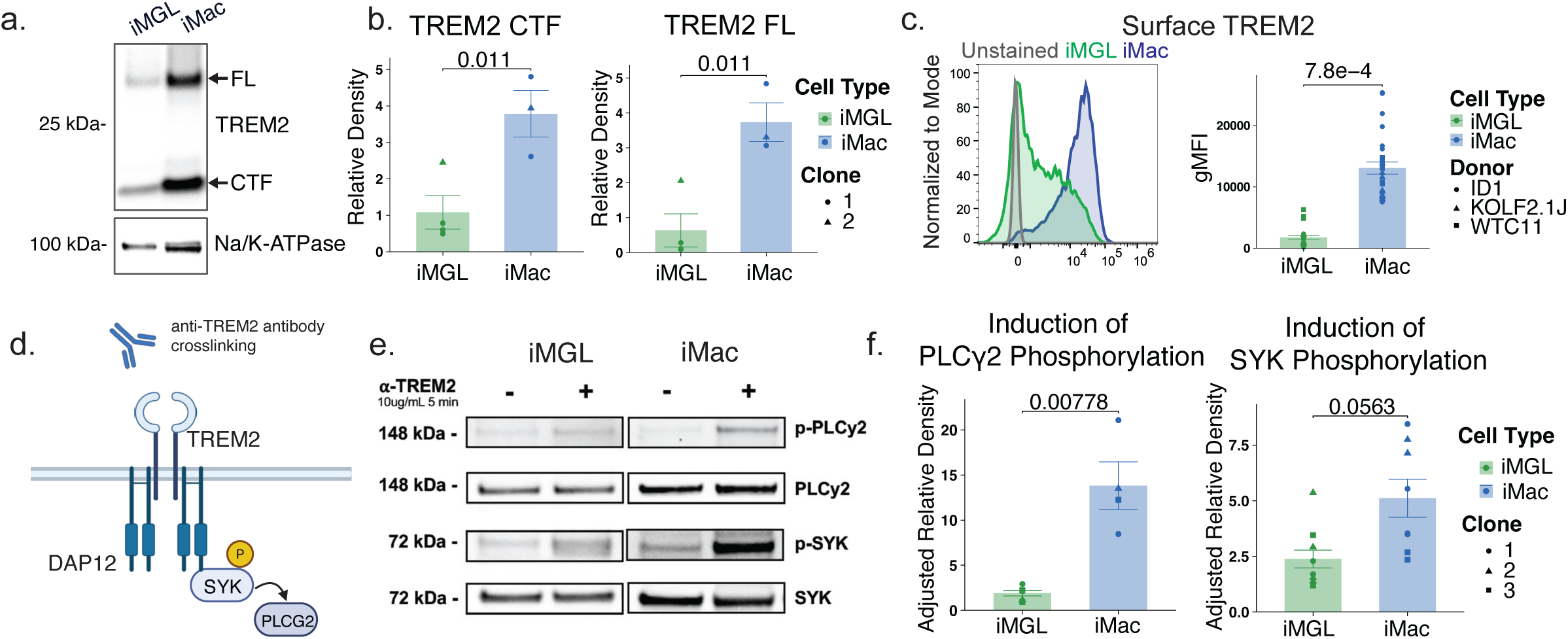
Increased TREM2 expression and downstream signaling activation in iMacs compared to iMGLs. **(a)** Representative western blot and **(b)** densitometry quantification of full-length TREM2 (TREM2-FL) and C-terminal fragment (TREM2-CTF) in the membrane fraction of WTC11 iMGLs and iMacs run side by side on the same blots, with Na/K-ATPase as loading control. **(c)** Representative flow cytometry histograms and surface expression quantification (gMFI) of TREM2 in iMGLs (green) and iMacs (blue) relative to unstained controls (gray). **(d)** Schematic of TREM2 antibody crosslinking and activation of the downstream SYK and PLCγ2 phosphorylation cascade. **(e)** Representative western blots of total and phosphorylated PLCγ2 and SYK following α-TREM2 treatment (10 µg/mL, 5 min) versus IgG-treated controls. **(f)** Quantification of fold induction of PLCγ2 and SYK phosphorylation upon TREM2 stimulation in iMGLs (green) and iMacs (blue). Experiments were conducted using 1–3 donor lines (1–3 clones per donor) across 2–3 independent differentiations per assay. Statistical significance was evaluated using linear mixed effects models. Full statistics are provided in the Supplementary Statistical Report.

We next sought to assess whether the elevated surface expression of TREM2 in iMacs translated to enhanced downstream signaling upon activation **(Fig. 4d-e)**. While antibody crosslinking of TREM2 induced rapid phosphorylation of downstream signaling proteins SYK and AD risk gene PLCγ2 in both cell types, iMacs demonstrated a stronger fold-induction of phosphorylated PLCγ2 (8.15-fold greater, 95% CI: 2.36–27.9, *P*_ad_ = 0.008) and phosphorylated SYK (1.71-fold greater, 95% CI: 0.98–2.97, *P*_ad_ = 0.056) relative to stimulated iMGLs **(Fig. 4e)**. These data establish that iMacs provide a highly responsive human cellular model for investigating TREM2-dependent signaling mechanisms.

### iMacs provide a highly efficient and reproducible system for siRNA-mediated gene perturbation

A major challenge with human pluripotent stem cell-derived microglia is their sensitivity to transfection reagents. Standard lipid-mediated transfection protocols frequently trigger cell toxicity in homeostatic iMGLs, while CRISPR/Cas9 knockout line generation is very expensive and time-consuming and can disrupt early myeloid differentiation if essential genes are deleted(Dräger et al. 2022). Moreover, the substantial cost and labor required to generate and validate independently edited lines often limit the number of genetically independent replicates that can be studied, reducing statistical power and leading to pseudoreplication when multiple measurements or differentiated cultures derived from the same edited clone are treated as independent biological replicates(Panda et al. 2023).

To determine whether iMacs provide a more robust platform for transient genetic manipulation, we evaluated siRNA-mediated knockdowns across multiple genetically diverse donor lines (ADRC5, ID1, KOLF2.1J, WTC11, and 051118). As a proof of concept, we targeted the AD-associated transcription factor *MEF2C* (**Fig. 5a**) which is predicted to regulate efferocytic gene programs such as lysosomal clearance and cholesterol efflux(Podlesny-Drabiniok et al. 2026). Transfection with MEF2C-targeted siRNA significantly reduced MEF2C protein levels (adjusted for actin) in both iMacs (72.0% reduction, 95% CI: 53.1–83.5%, *P*_adj_ = 2.5 × 10^−4^) and iMGLs (44.4% reduction, 95% CI: 23.0–59.6%, *P*_adj_ = 0.002) (**Fig. 5b**). However, knockdown was significantly more efficient in iMacs (genotype × cell type interaction, *P* = 0.042, **Fig. 5c**) and and more consistent across donors (57.6–83.4% reduction across iMac donors, significant in 3 of 4, compared with 10.2–67.9% and 1 of 4 across iMGL donors, **Supplementary Statistical Reporting Document**), supporting iMacs as a reproducible and efficient system for siRNA-mediated gene perturbation across genetically diverse donors.

**Fig 5.**
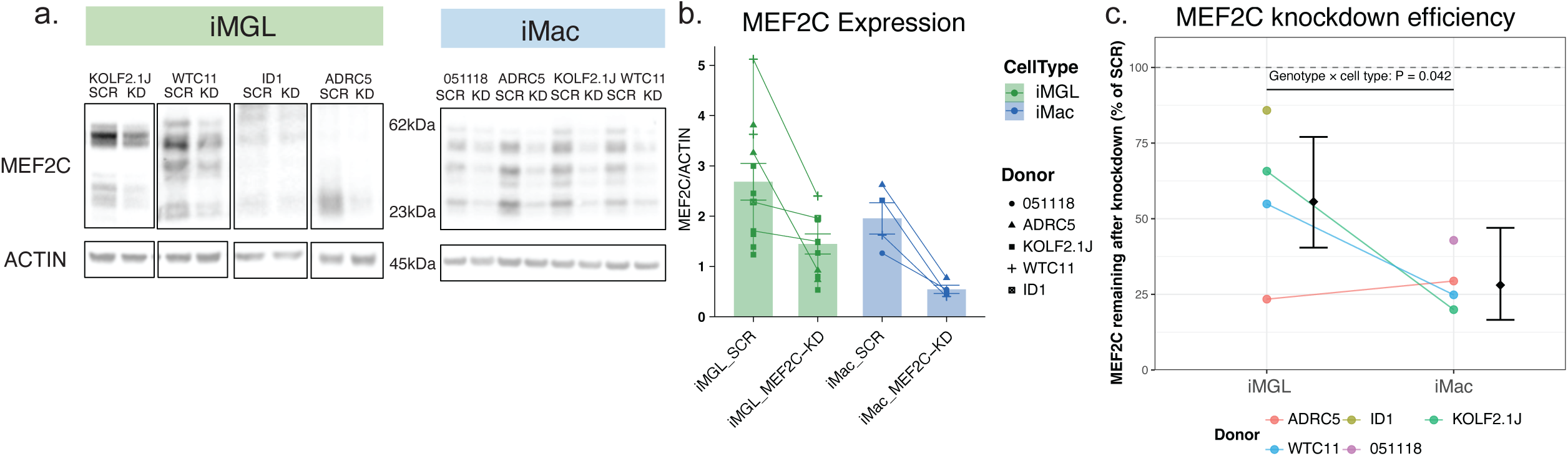
Efficient siRNA-mediated gene knockdown in iMacs compared to iMGLs across multiple iPSC donor lines. (a) Representative western blot analysis of MEF2C protein expression following siRNA knockdown (KD) compared to scrambled non-targeting control (SCR) in iMGLs and iMacs across multiple distinct iPSC donor lines. ACTIN serves as a loading control. Molecular weight markers are indicated in kilodaltons (kDa). **(b)** Densitometric quantification of MEF2C relative to ACTIN in iMGLs (green) and iMacs (blue) under SCR and MEF2C-KD conditions. Small points represent individual lanes and large symbols represent donor means, with shapes indicating distinct iPSC donor lines (4 donors per cell type; 5 lines in total). Lines connect SCR and KD conditions within each donor; bars show mean ± SEM of donor means. *P*_adj_ values indicate SCR vs. KD within each cell type. **(c)** MEF2C knockdown efficiency, expressed as MEF2C remaining after KD (% of SCR). Points show individual donors, with lines connecting the same donor across cell types; diamonds and error bars show model-estimated % MEF2C remaining with 95% confidence intervals. *P* value indicates the genotype × cell type interaction. Statistical significance was evaluated using linear mixed effects models. Full statistics are provided in the Supplementary Statistical Report.

## Discussion

In this study, we describe a rapid, 19-day protocol for deriving functional human macrophage-like cells (iMacs) from iPSC-derived EMP-like HPC intermediates using M-CSF as a tractable system for studying efferocytosis in a disease-relevant human cell type. The recognition, uptake, and lysosomal degradation of extracellular substrates are functions repeatedly implicated in Alzheimer’s disease by human genetics studies. Compared with iMGLs, iMacs show enhanced capacity for these processes both at the gene expression and functional level. By using a cryopreservable Day 12 HPC intermediate and a simplified maturation phase, this workflow avoids the lengthy schedules, costly cytokine supplementation, and viral transduction required by current microglial culture protocols. A key strength of our iMac protocol is its practical utility. Roughly halving the differentiation time reduces media and growth factor costs, while yielding highly pure (>90%), viable cultures with high batch-to-batch consistency across diverse donor lines. Moreover, because iMacs derive from EMP-like HPCs rather than monocytes, they retain a developmental origin that more closely resembles that of TRMs such as microglia and BAMs.

Functionally, iMacs express high baseline levels of the receptors that initiate efferocytosis in tissue-resident macrophages, including the key BAM identity markers CD36, CD163 and CD206. They also express high levels of phagocytic receptors such as CD64, TREM2, and Dectin-1 (CLEC7A), with TREM2 and Dectin-1 being of particular interest due to their role as canonical DAM markers. Together with an expanded lysosomal compartment, these features most likely promote the rapid internalisation of diverse substrates, including latex beads, zymosan, Aβ 1-42, and Aβ 1-40. This is particularly relevant to AD, where both TREM2 and CD36 mediate the phagocytosis of amyloid-β(S.-M. Kim et al. 2017; El Khoury et al. 2003) and where TREM2 is itself a well-established risk gene. Consistent with this, TREM2 engagement elicited markedly greater p-SYK and p-PLCγ2 responses in iMacs than in iMGLs. In line with the convergence of AD genetic risk on macrophage efferocytosis, the iMac transcriptional program was selectively enriched for AD risk genes relative to those of other neurodegenerative diseases(Romero-Molina et al. 2022). Notably, iMacs acquire these features at baseline and do not require the substrate exposure, transcription factor overexpression, or engineered knockout lines needed to induce DAM-like states in iMGL systems(Dolan et al. 2023; Podleśny-Drabiniok et al. 2025; Podlesny-Drabiniok et al. 2026).

Transcriptomic benchmarking placed iMacs closest to CNS macrophages among the tissue-resident populations examined. Within that compartment they were enriched for non-homeostatic states, including BAM and DAM signatures. Since our protocol omits the brain parenchymal signals used in iMGL differentiation, iMacs may be prevented from acquiring the homeostatic microglial transcriptome. This could position iMacs as a human model of CNS macrophage states most closely associated with neurodegeneration. However, this identity does not appear to be exclusively CNS-associated. The enrichment of LAM signatures in iMacs reflects the lipid-rich nature of the brain, a shared trait with peripheral organs such as the liver in which the receptor and clearance programs of LAMs were first described. iMacs may therefore be useful for studying disease-associated macrophages in lipid-rich organs beyond the brain, including those in atherosclerosis, fatty liver disease, and obesity.

A further advantage of this system is its suitability for acute genetic manipulation by lipid-mediated transient transfection. Lipid-mediated transfection yields lower knockdown efficiency in iMGLs, whereas iMacs reliably achieve strong siRNA knockdown of target genes such as MEF2C across multiple donor lines. This allows researchers to manipulate candidate genes and assess their effects on macrophage function within days, rather than the months needed to generate and validate genome-edited lines. Combined with the speed of the differentiation itself, this makes iMacs well suited to screening approaches in which many genes or compounds must be tested in parallel. In conclusion, our iMac protocol provides a fast, economical, and robust human macrophage model that bridges the gap between complex microglial differentiation protocols and the practical demands of studying disease-relevant macrophage function at scale.

### Limitations & Future Directions

This protocol lacks the cell-cell contacts, extracellular matrix, and three-dimensional architecture of the brain, and omits brain parenchymal signals such as TGF-β and IL-34. As such, the states iMacs adopt are shaped by culture conditions rather than by a specific tissue environment. Accordingly, while iMacs share key features with CNS macrophages such as BAM and DAM, the resemblance is partial and cannot be assumed to be equivalent to macrophage identities continuously reinforced by the specific local cues found *in vivo*. Because all transcriptomic profiling was performed by bulk RNA sequencing, cellular heterogeneity within iMac cultures remains uncharacterized. Single-cell approaches would be required to determine whether the BAM- and DAM-associated signatures reflect a uniform population or distinct subpopulations. Their efferocytic phenotype was also characterised using a limited set of substrates, which should be explored further by testing additional material such as myelin and apoptotic cells. Genetic tractability has thus far been demonstrated for a single target gene. Though their amenability to genetic manipulation is promising, the suitability of iMacs for larger-scale perturbation such as a pooled CRISPR screen remains to be established. Future work could include re-introducing specific environmental factors to test whether iMacs can be selectively directed toward distinct tissue-resident macrophage subsets. As a tractable HPC-derived macrophage model, future exploration could also involve integration of iMacs into more complex systems such as co-cultures, organoids, and *in vivo* xenotransplantation models, to probe their interactions with other CNS cell types in health and disease.

## Methods

### Induced pluripotent stem cell (iPSC) lines

iPSCs from five donors (KOLF2.1J, WTC11, ID1, ADRC5, and x05118) were used in this study. The KOLF2.1J (Product Code “JIPSC001000”) iPSC line was provided by The Jackson Laboratory (JAX) as part of the NIH-sponsored iPSC Neurodegenerative Disease Initiative (iNDI) Differentiation Study. The remaining donor iPSC lines (WTC11, ID1, ADRC5, and 05118) were previously generated, characterized, and validated as described (Kreitzer et al. 2013; Tcw et al. 2022; Podlesny-Drabiniok et al. 2026; Weitzman et al. 2026). All iPSC lines were maintained using StemFlex Medium (Thermo Fisher Scientific) and routinely tested for mycoplasma using the MycoStrip Mycoplasma Detection Kit (Invivogen, catalog #rep-mysnc-100) to prevent potential contamination or alteration to the cell lines.

### Differentiation of iMGLs

iPSC-derived microglia-like cells (iMGLs) were differentiated following a previously published paper (Abud and McQuade) with minor modifications. iPSCs were passaged every 5–6 days using ReLeSR dissociation reagent (STEMCELL Technologies, cat. 05872) and used for hematopoietic stem cell differentiation with STEMdiff Hematopoietic kit (STEMCELL Technologies, cat. 05310) to generate hematopoietic precursor cells (HPCs). Cells were then replated in iMGL media supplemented with 3 cytokines (100 ng/ml IL34, 50 ng/ml TGFβ, 25 ng/ml MCSF) on Matrigel coated plates for 25 days, and additionally supplemented with 2 cytokines (CX3CL1 and CD200, 50 ng/ml each) for an additional three days. Mature iMGLs (day 28) were used for all downstream experiments.

### Differentiation of iMacs

We adapted a published STEMCELL Technologies protocol (Tabatabaei-Zavareh et al, 2017) to differentiate iMacs from HPCs instead of monocytes to better mimic EMP-derived tissue resident macrophages. HPCs were cultured in macrophage medium (STEMCELL Technologies, 10961) supplemented with 50 ng/ml MCSF for a minimum of 6 days. 24 hours prior to harvest, cells were lifted with PBS (Gibco 14190-144) and plated in fresh media for downstream analyses.

### RNAseq Analysis

#### Differential Expression, Pathway and Hypergeometric Overlap Testing

FastQ files for samples were first processed through the Nextflow RNA-Seq pipeline. This included assessment of sequencing read quality (FastQC), adapter contamination trimming (Cutadapt), alignment of reads to the human genome hg38 (STAR), and quantification of counts at gene level with SALMON. Differential expression analysis was then performed in DREAM (variancePartition package), employing a mixed-effects model with “Cell_Type” and “Clone” specified as fixed and random effects respectively, with p-values adjusted by Benjamini-Hochberg correction. Differentially expressed genes were then ranked by t-statistic, and gene-set enrichment analyses (GSEA) were carried out with the fgsea package, against genesets from the Molecular Signatures Database (MSigDB v7.5.1). GSEA against published microglial and macrophage state signatures, including disease-associated microglia, was performed with the clusterProfiler package, using gene signatures compiled in Podlesny-Drabiniok et al. (2026). Each GSEA was corrected separately by Benjamini-Hochberg within its own gene set collection.

Enrichment of disease risk genes among differentially expressed genes was assessed by one-sided hypergeometric test (phyper, R stats package). Risk gene lists were obtained from the NHGRI-EBI GWAS Catalog (https://www.ebi.ac.uk/gwas/) for Alzheimer’s disease (AD; Jansen et al. 2019; Kunkle et al. 2019; Bellenguez et al. 2022), Parkinson’s disease (PD; Nalls et al. 2019; Kim et al. 2024), amyotrophic lateral sclerosis (ALS; van Rheenen et al. 2021) and multiple sclerosis (MS; International Multiple Sclerosis Genetics Consortium 2019), using the Catalog’s mapped genes (the gene overlapping the lead variant or, for intergenic variants, the nearest upstream and downstream genes). For each disease, the overlap between its risk genes and all genes differentially expressed between iMacs and iMGLs (FDR < 0.05 and |log₂FC| > 1, up- and downregulated genes tested together; 4,136 genes) was evaluated against a universe comprising all genes tested in the differential expression analysis (14,587 genes). Duplicated gene symbols were collapsed to the most significant entry, and risk genes absent from the tested universe were excluded from the list size rather than counted as non-overlapping. Fold enrichment was calculated as the overlap rate within each risk gene list divided by the transcriptome-wide DEG rate. Odds ratios and two-sided exact 95% confidence intervals were obtained by Fisher’s exact test (fisher.test), and p-values were adjusted across the four diseases by Benjamini–Hochberg correction. To account for the greater statistical power to detect differential expression among highly expressed genes, an expression-matched permutation null was also generated by resampling each risk gene list 10,000 times from the universe, matched on expression decile, and the observed overlap was compared against this null distribution.

#### Principal Component Analyses

FastQ files from publicly available studies with cell types of interest for PCA were downloaded from NCBI SRA selector or synapse data repositories. Those from bulk-RNAseq datasets were then processed through the Nextflow RNA-seq pipeline detailed earlier. FastQ files from scRNAseq datasets (all generated with 10X Genomics technology) were processed through CellRanger for mapping of reads to reference genomes (GRCh38) and generation of feature and count matrices for downstream analysis in Seurat to identify and annotate microglia and border-associated macrophages. Ambient RNA removal was also performed prior to analysis in Seurat with CellBender for FreshMG cohort data from (Lee et al. 2026). Following loading of feature-barcode matrices into Seurat, doublets were identified with scDblFinder and removed, along with cells that had a mitochondrial genome content >10-15% (threshold determined for each individual study after inspection of histograms for mitochondrial content measures). Normalization and batch correction were performed with SCTransform (with mitochondrial content regressed out where needed) and Harmony packages respectively. Clusters were identified with Seurat’s FindClusters() function following identification of the 2,000 most variable features in each dataset. Clusters were annotated following differential expression analysis with the presto package, and macrophage clusters were identified by expression of canonical markers such as PTPRC, AIF1, CSF1R, C1QA, ITGAM, and absence of markers of other major cell types. PCA was performed on log-normalised count matrices using prcomp() from the stats package, with features scaled to unit variance. For the comparison against primary tissue-resident macrophages, all genes shared across cohorts were used, excluding only those with zero variance; for the comparison against *in vitro* myeloid models, the 2,000 most variable features were used. For the tissue-resident macrophage comparison, batch correction was applied prior to prcomp() using removeBatchEffect() from the limma package, with assay platform (bulk RNA-seq versus pseudobulk derived from scRNA-seq) as the batch variable. Group dispersion was visualised as 95% normal data ellipses, computed from the covariance of the PC1 and PC2 scores assuming a bivariate normal distribution (stat_ellipse(type = “norm”, level = 0.95) in ggplot2); ellipses were not drawn for groups with fewer than three samples. Similarity between iMacs and other cell types was quantified using Euclidean distances and Kullback–Leibler (KL) divergence. Euclidean distances were computed between each sample and the iMac centroid using sample loadings on PC1 and PC2. KL divergence was calculated in base R from the closed-form expression for two multivariate normal distributions, comparing each group’s bivariate Gaussian approximation in PC1/PC2 space with that of the iMac reference group; groups with fewer than two samples were excluded.

### Lysosomal Assays

To measure lysosomal mass and acidification, cells were incubated with 75 nM LysoTracker-Red (Thermo Fisher Scientific, L7525) for 30 min at 37°C followed by 1 µM LysoSensor-Green (Thermo Fisher Scientific, L7535) for 1 min at 37°C. Cells were analyzed via flow cytometry using an Attune cytometer (Thermo Fisher Scientific) and analyzed using FlowJo Analysis Software.

### Latex bead Uptake

Uptake of fluorescent beads was measured by flow cytometry. iMGLs and iMacs were plated 30,000 cells/well in a 96 well matrigel-coated (Corning, cat. 354230) plate. Cells were treated with GFP-labeled carboxylate-modified polystyrene 2 micron beads (Sigma-Aldrich, cat. L4530-1ML) for 24 hours. To inhibit uptake, cells were pre-treated with 2 μM Cytochalasin D (Sigma-Aldrich, cat. C8273-1MG) for 30 min and during incubation with bioparticles. After 24-hour incubation cells were collected with Accutase (Stemcell Technologies, cat. 07920) for 10 min at 37°C and resuspended in 1.0% PBS/bovine serum albumin (BSA) buffer and analyzed on an Attune flow cytometer (Thermo Fisher Scientific). Data were analyzed using FCS Express 7 (De Novo Software). Following doublet exclusion, fluorescence intensity was used to quantify bead number within each cell. gMFI and bead numbers were analyzed and plotted.

### Amyloid and Zymosan Phagocytosis Live Imaging assays

For phagocytosis experiments, cells were plated at a density of 30,000 cells/well and treated with either 1 µM 488-conjugated amyloid 1-42 (Anaspec, AS-60479-01), 1 µM 488-conjugated amyloid 1-40 (Anaspec, AS-60491-01) or 5 ug/mL pHrodo-Zymosan (Thermo Scientific, P35364). As a control, cells were pre-treated with 2uM cytochalasin-D for 30 minutes prior to addition of substrates. Cell confluence and red/green fluorescence signal were quantified for 24h using the Incucyte S3 live imaging system. Total integrated density was calculated as mean fluorescence intensity multiplied by surface area of masked object [RCU x µm2] and normalized by cell confluence (phase channel).

### TREM2 Signaling Assays

Cells plated in a 6 well plate at a density of 500,000 cells per well were lifted with PBS and spun down at 300g for 5 minutes. Cell pellets were resuspended in HBSS containing either polyclonal goat anti-TREM2 (R&D Systems, AF18281) or polyclonal goat isotype control (R&D Systems, AB-108-C) at 10 µg/ml. Cells were incubated at 37°C in a heat block for 5 minutes. Reactions were quenched with ice cold PBS containing Protease/Phosphatase Inhibitor Cocktail (Cell Signaling, 5872). Cells were spun at 300g for 5 minutes at 4°C, supernatant removed, and either frozen or directly processed for western-blotting.

### siRNA Transfection

Mature iPSC-derived microglia (iMGLs; day 28 of differentiation) and iPSC-derived macrophages (iMacs; day 6 of M-CSF–mediated differentiation) were transfected with 40 nM ON-TARGETplus Human MEF2C siRNA SMARTpool (Horizon Discovery, L-009455-00) or 40 nM ON-TARGETplus Non-targeting Control siRNA (Horizon Discovery, D-001810-10-05). Transfections were performed using Lipofectamine RNAiMAX (Thermo Fisher Scientific, LMRNA015) according to the manufacturer’s instructions. Cells were maintained in their respective culture media and harvested 48 h after transfection for downstream RNA and protein analyses. MEF2C knockdown was validated by immunoblotting, with MEF2C protein intensity normalized to Actin. Experiments were performed across multiple independent iPSC donors and, where available, multiple differentiations per donor.

### Western-blotting

For total protein measurements, cells were lysed in RIPA buffer (Thermo Scientific, 89900) supplemented with Protease/Phosphatase Inhibitor Cocktail (Cell Signaling, 5872) following manufacturer’s instructions. Lysed cells were sonicated and spun down to pellet debris prior to freezing supernatants. For membrane protein measurements, samples were lysed using the MemPer Protein Extraction kit (Thermo Scientific, 89842) supplemented with Protease/Phosphatase inhibitor cocktail to enrich membrane-bound proteins separately. Protein concentration was measured using BCA kit (Thermo Fisher Scientific, 23225) and equal quantities (10-20ng) were used to prepare samples for western blotting. Samples were resolved by electrophoresis with Bolt 4-12% Bis-Tris Plus Gels (Invitrogen) in Bolt MES SDS running buffer (Invitrogen, B0002) and transferred using iBlot 3 nitrocellulose transfer stacks (Invitrogen) using program P0. Membranes were blocked for 1 h and probed with primary antibodies in 5% BSA in PBS/0.1% Tween-20 buffer overnight at 4°C (**Table 1)**. Secondary antibody at 1:10,000 dilution was applied for 1 h at RT, and visualized using WesternBright ECL HRP Substrate Kit (Advansta, K-12045) or SuperSignal™ West Femto Maximum Sensitivity Substrate (Thermo Scientific, 34094), and measured using iBright imaging system (Applied Bioscience). Images were analyzed using ImageJ.

**Table 1:** Primary antibodies for western-blotting.

| **Antibody** | **Reactivity** | **Host** | **Manufacturer** | **Catalog** | **Concentration** |
| --- | --- | --- | --- | --- | --- |
| *Na,K-ATPase* | Human | Rabbit | Cell Signaling Technology | 23565S | 1:200 |
| *P-PLCg2 (Tyr1217)* | Human | Rabbit | Cell Signaling Technology | 54442T | 1:500 |
| *PLCg2* | Human | Rabbit | Cell Signaling Technology | 55512 | 1:500 |
| *P-SYK (Tyr352)* | Human | Rabbit | Cell Signaling Technology | 2717 | 1:500 |
| *SYK* | Human | Rabbit | Cell Signaling Technology | 13198 | 1:500 |
| *TREM2* | Human | Rabbit | Cell Signaling Technology | 91068 | 1:500 |
| *MEF2C* | Human | Rabbit | abcam | 211493 | 1:1000 |
| *ACTIN* | Human | Mouse | abcam | 7817 | 1:10,000 |

### Flow Cytometry

Cells were harvested at 400 × g for 5 min, and resuspended in cold D-PBS containing 1.0% bovine serum albumin (BSA). To assess viability and prevent non-specific binding, cells were incubated with Live/Dead Fixable Violet Dead Cell Stain (Life Technologies L23105) and Human Fc Block (BD Biosciences, cat. 564220) in D-PBS for 30 min at 4°C. Following two PBS washes, cells were stained with fluorophore-conjugated antibody cocktails designed to evaluate cell identity and DAM signatures. Staining was performed in a 1:1 mixture of FACS buffer (D-PBS + 0.5% BSA) and Brilliant Stain Buffer (BD Biosciences, cat. 563794) for 30 min at 4°C. Markers analyzed in this study included surface targets (CD206-BUV395 (1:25, BD Biosciences, cat. 740309), HLA-DR-BV605 (1:100, BioLegend, cat. 307640), CD11b-BV785 (1:100, BioLegend, cat. 101243), CD45-PerCP (1:50, BD Biosciences, cat. 368506), CD163-PECy5 (1:100, BioLegend, cat. 333644), CD64-APC (1:100, BioLegend, cat. 305014), Clec7a-PerCP-eFluor 710 (1:100, Thermo Fisher Scientific, cat. 46-9856-42), CD36-PE-Fire 640 (1:100, BioLegend, cat. 336246), TREM2-APC (1:50, R&D Systems, cat. FAB17291A) and intracellular targets (CD68-BV421 (1:50, BD Biosciences, cat. 333828) and LAMP1-BV510 (1:100, BD Biosciences, cat. 328632). For intracellular staining, cells were fixed and permeabilized using the Foxp3 Fixation/Permeabilization Buffer Set (Thermo Fisher Scientific, cat. 00-5523-00) for 30 min at room temperature, followed by incubation with intracellular antibodies for 60 min at 4°C in 1× Permeabilization Buffer. Cells were washed twice in FACS buffer and acquired using a 5-laser Cytek Aurora spectral flow cytometer (Cytek Biosciences). Single-stain reference controls were prepared using ultra-comp capture beads (Invitrogen, cat. 01-3333-42) and cellular controls. Data analysis was conducted using FlowJo software v10.8 (BD Biosciences), and target expression levels were quantified as geometric mean fluorescence intensity (gMFI).

### Statistical analysis

All statistical analyses were carried out in R, using linear-mixed effects modelling (‘lme4’ package). Models were first assessed for linearity, homogeneity of variance and normality with the performance package to ensure model assumptions were met. If assumptions were not satisfied, data was either transformed, or a generalised mixed effect linear model with an appropriate family suited to the distribution of the data was used to test for significance. For datasets with multiple continuous dependent variables examined simultaneously, data were standardized using the scale() function in R. The estimate_contrasts() function of the modelbased package was employed to calculate estimated marginal means, confidence intervals and adjusted p-values (through ‘bonferroni’ or ‘fdr’ correction) for each contrast. Where data were transformed, *P* values were obtained on the transformed scale, and estimated marginal means were back-transformed to the original scale for reporting; effect sizes (β) are reported as differences in back-transformed estimated marginal means with unadjusted 95% confidence intervals.

## Supporting information

Supplementary Statistical Reporting Document

## Data and Code availability

Fastq files of external datasets analyzed in this manuscript are available to download from NCBI GEO for Tuddenham et al., (GSE204702), Abud et al., (GSE89189), Brownjohn et al., (GSE110952), Drager et al., (GSE178317), Liu et al., (GSE287851), Podlesny-Drabinoik et al., (GSE273482), Quiroga et al., (GSE181153), Sauer et al., (GSE298244), Zhang et al., 2025 (GSE55536), Penney et al., (GSE241858), Sun et al., (GSE227223), Ramaswami et al., (GSE226081), McQuade et al., (GSE158469), Haskell et al., (GSE306820) and Hasselman et al., (GSE133432), and from the Synapse portal for Lee et al., (syn52795287). Raw data and code generated from this study will be deposited onto Github and GEO Omnibus upon publication. Requests for access to raw data and analysis scripts should be directed to the corresponding author.

## Acknowledgements

**<u>Grants</u>** This work was funded by the following grants: NIH U19AG069701 (A.M.G.), R01HL168174 (E.M.), R01AG072489 (A.M.G.), F32AG092137 (H.L.), F31AG084337 (A.M.), T32AI078892 (J.J.K.), Cure Alzheimer’s Fund (H.L., A.M.G.), BrightFocus Foundation A2024025S (A.P-D.), The Freedom Together Foundation (A.M.G.), and The Belfer Neurodegeneration Consortium (A.M.G., E.M., M.S., G.B.P). A.M.G and E.M acknowledge funding from the Sanford J Grossman Charitable Trust. **<u>Cores:</u>** Research reported in this publication was supported by the Flow Cytometry CoRE (RRID:SCR_027701) at the Icahn School of Medicine at Mount Sinai and by the National Cancer Institute of the National Institutes of Health under award number P30 CA196521. Experiments were supported by the Stem Cell Engineering Core (RRID:SCR_027503) at the Icahn School of Medicine at Mount Sinai. This work was supported in part through the computational and data resources and staff expertise provided by Scientific Computing and Data at the Icahn School of Medicine at Mount Sinai and supported by the Clinical and Translational Science Awards (CTSA) grant UL1TR004419 from the National Center for Advancing Translational Sciences. Research reported in this publication was also supported by the Office of Research Infrastructure of the National Institutes of Health under award number S10OD026880 and S10OD030463. The content is solely the responsibility of the authors and does not necessarily represent the official views of the National Institutes of Health. Figure 1a, 5d, and Supplementary Figure 5a were created in https://BioRender.com **<u>iPSC line:</u>** We thank the NYSCF, Mount Sinai Stem Cell Core, Washington University in St. Louis Knight ADRC (P30AG066444), and University of California, Irvine ADRC (P30AG066519) for providing the iPSC lines for this project. KOLF IPSC line: Product was generated from the parental iPS cell line produced by Genome Research Limited, operating as the Wellcome Trust Sanger Institute, as part of the Human Induced Pluripotent Stem Cell Initiative (HIPSCI) and was generated with the support of the National Institutes of Health (NIH) and the NIH’s iPSC Neurodegenerative Disease Initiative.

## Author contributions

Conceptualization, H.L., G.B.P., M.S., E.M., and A.M.G.; methodology, H.L., G.B.P., M.S., J.O., J.K., A.P.D., and A.M.; software, M.S., and E.M.; validation, H.L., G.B. P., M.S., A.P.D., and A.M.; formal analysis, H.L., G.B.P., and M.S.; investigation, H.L., G.B.P., M.S., J.O., J.K., A.P.D., and A.M.; resources, E.M., and A.M.G.; data curation, H.L., G.B.P., and M.S.; writing – original draft, H.L., G.B.P., and M.S.; writing – review & editing, H.L., G.B.P., M.S., J.O., J.K., A.P.D., A.M., E.M., and A.M.G.; visualization, H.L., G.B.P., and M.S.; supervision, E.M., and A.M.G.; project administration, E.M., and A.M.G.; funding acquisition, E.M., and A.M.G.

---

**Supplementary Figure 1.**
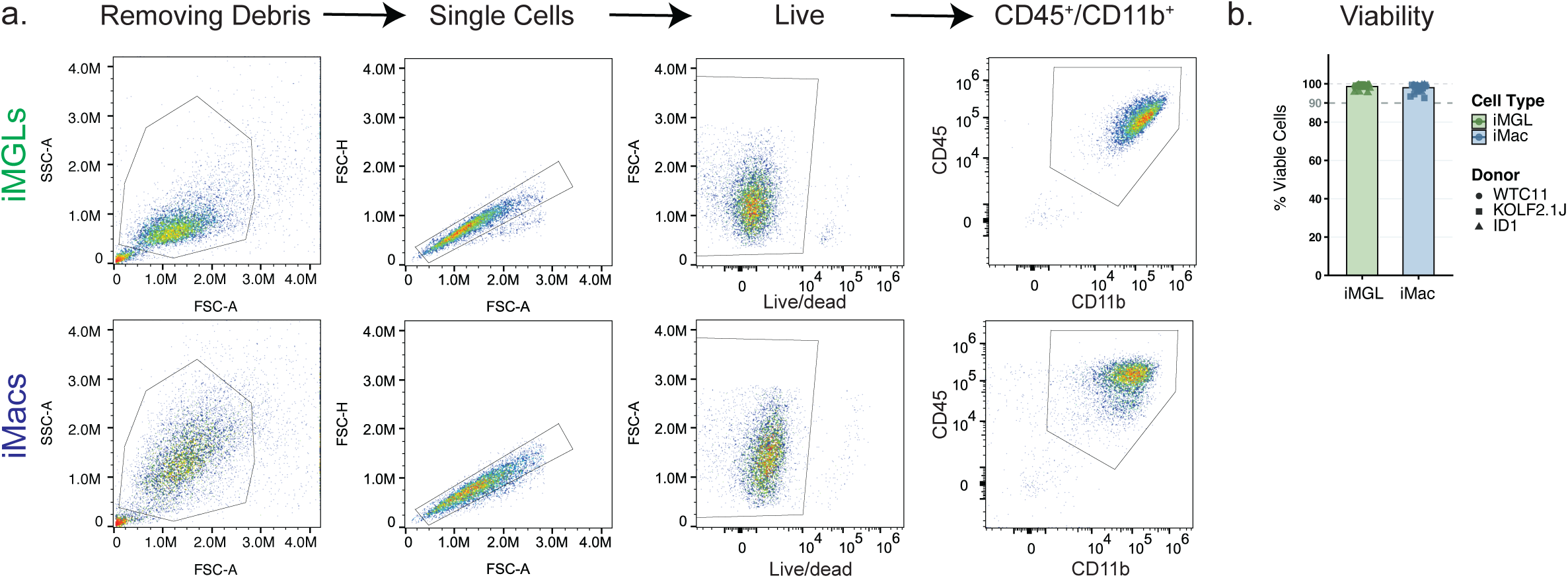
Characterization of myeloid differentiation efficiency. **(a)** Representative flow cytometry gating strategy for iMGLs (top) and iMacs (bottom). Sequential gating strategy identifies singlet cells (FSC-A vs. FSC-H), live/viable cells, and double-positive CD45⁺ CD11b⁺ myeloid populations. **b,** Quantification of cell viability (% viable cells) comparing iMGLs and iMacs across 3 differentiations, 3 iPSC donor lines. Dashed horizontal lines indicate the >90% purity threshold.

**Supplementary Figure 2.**
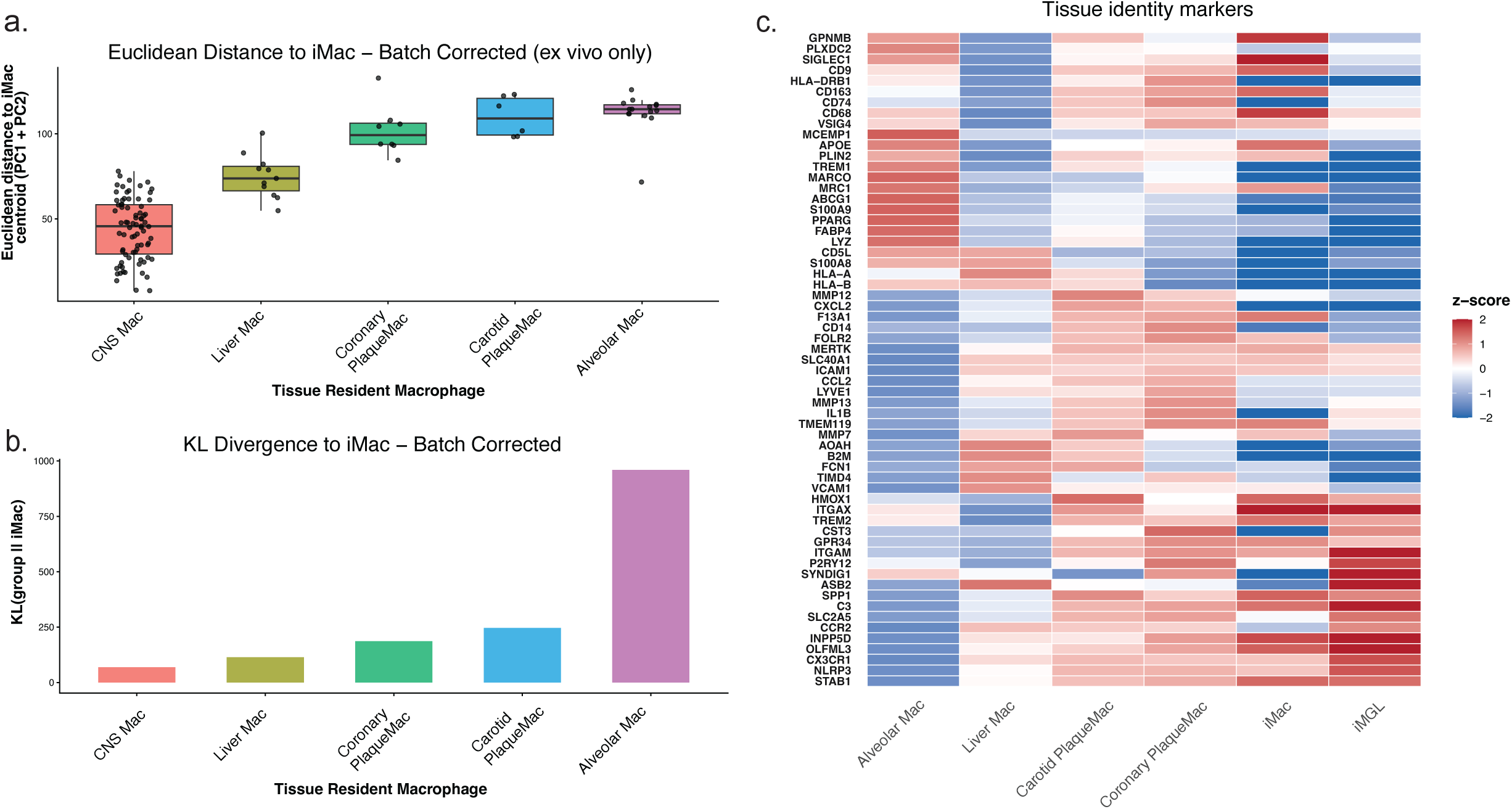
Transcriptomic comparison of iMacs and other tissue resident macrophage signatures. **(a)** Euclidean distance quantification and (b) (KL) divergence analysis measuring transcriptomic similarity between in-house iMacs and *ex vivo* tissue-resident macrophage lineages. (c) Heatmap showing relative expression (z score) of key tissue-identity marker genes that distinguish distinct human tissue-resident macrophage populations (alveolar, liver, atheroma, coronary) alongside iMGLs and iMacs.

**Supplementary Figure 3.**
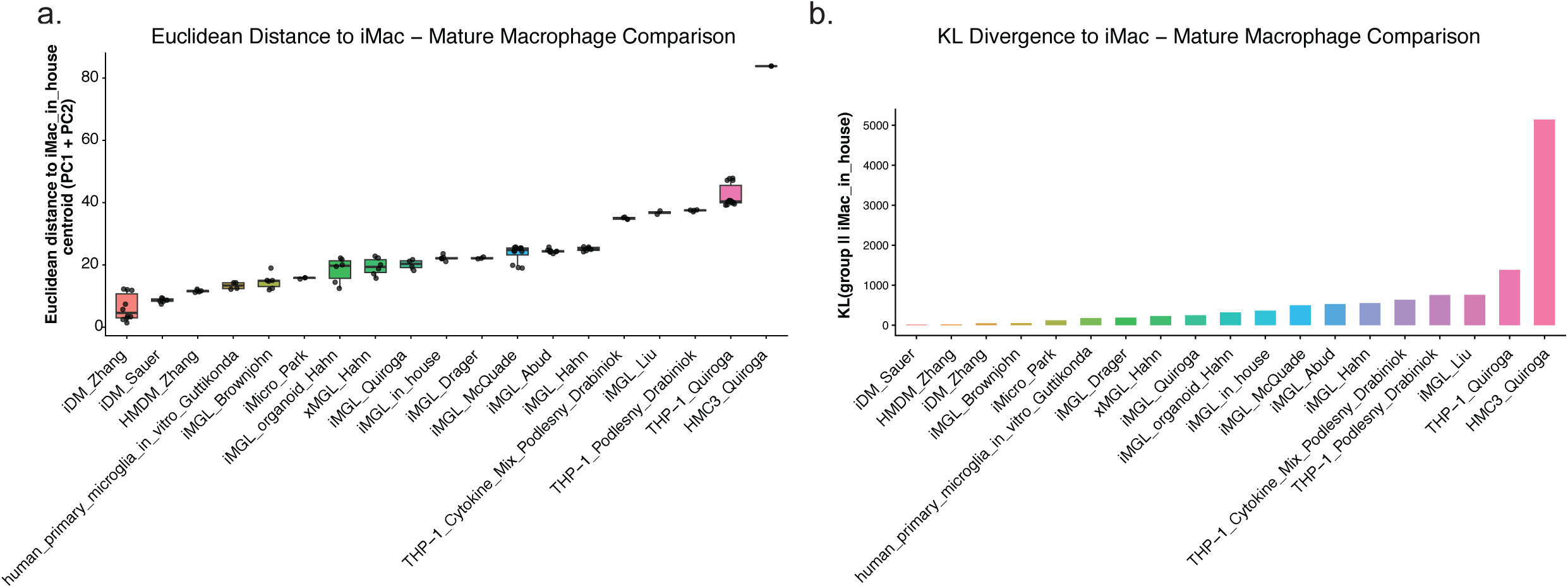
Transcriptomic comparison of iMacs and other mature macrophage dataset profiles. **(a)** Euclidean distance quantification across principal component space (PC1 + PC2) calculating relative similarity to in-house iMacs across distinct mature macrophage dataset profiles. **(b)** Kullback–Leibler (KL) divergence analysis relative to iMac expression profiles across mature macrophage subtypes.

**Supplementary Figure 4.**
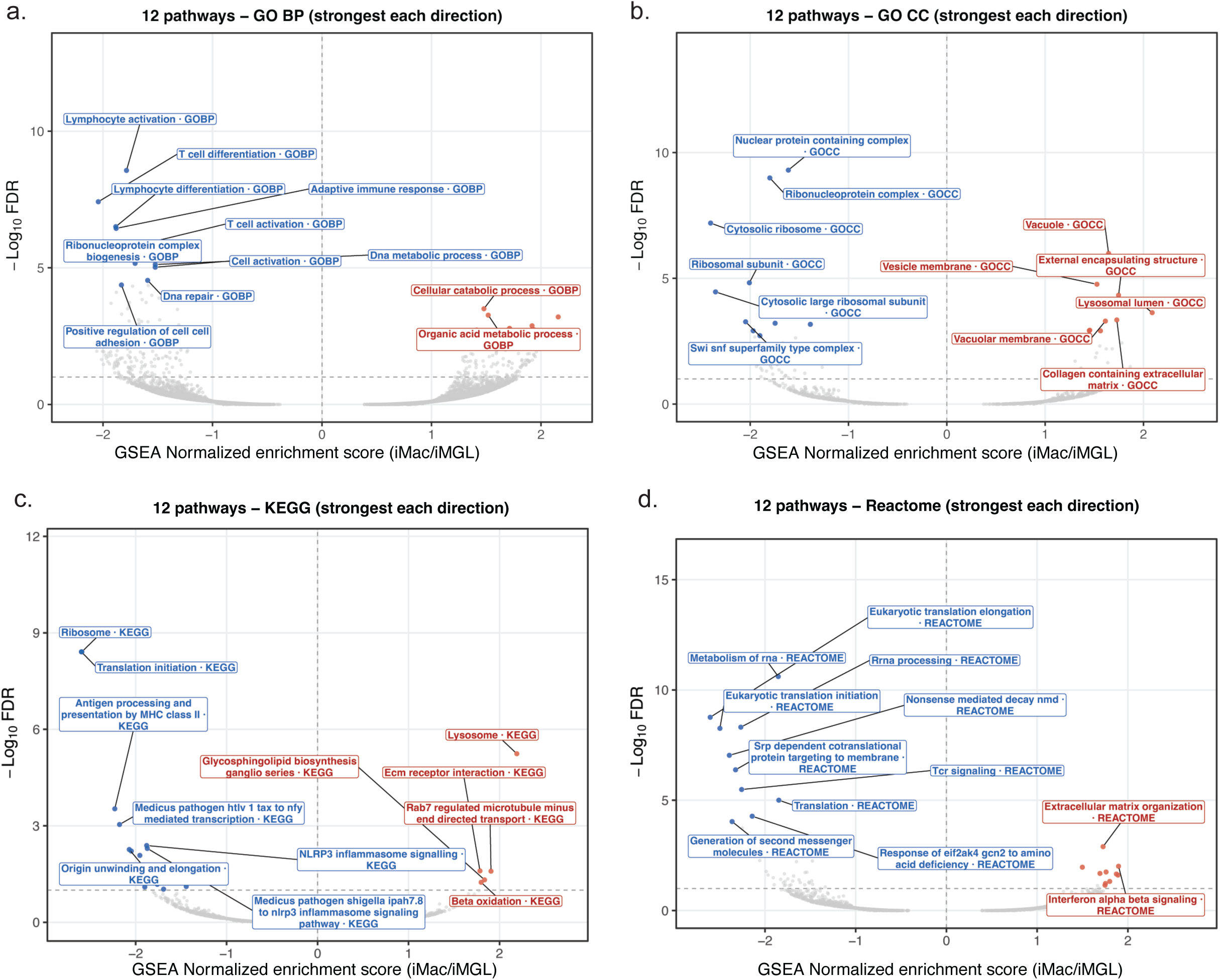
Unbiased GSEA in iMGLs vs. iMacs. (a–d) Volcano plots displaying the top enriched biological pathways in iMacs (red) versus standard iMGLs (blue) across curated pathway databases: **(a)** Gene Ontology Biological Process (GO BP), **(b)** Gene Ontology Cellular Component (GO CC), **(c)** KEGG pathways, and **(d)** Reactome pathways.

**Supplementary Figure 5.**
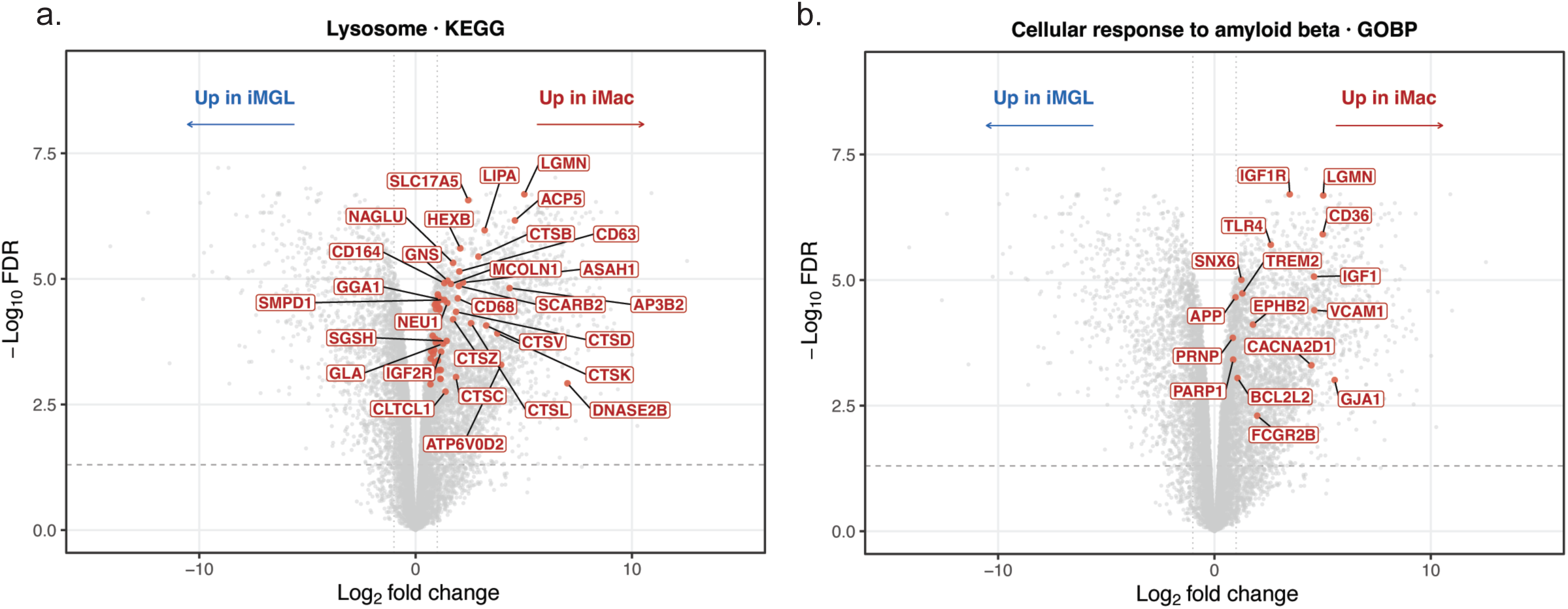
Leading-edge gene expression in iMGLs vs. iMacs. (a-b) Volcano plot highlighting leading-edge genes driving the pathways (a) KEGG Lysosome and (b) GO BP Cellular Response to Amyloid Beta.

**Supplementary Figure 6.**
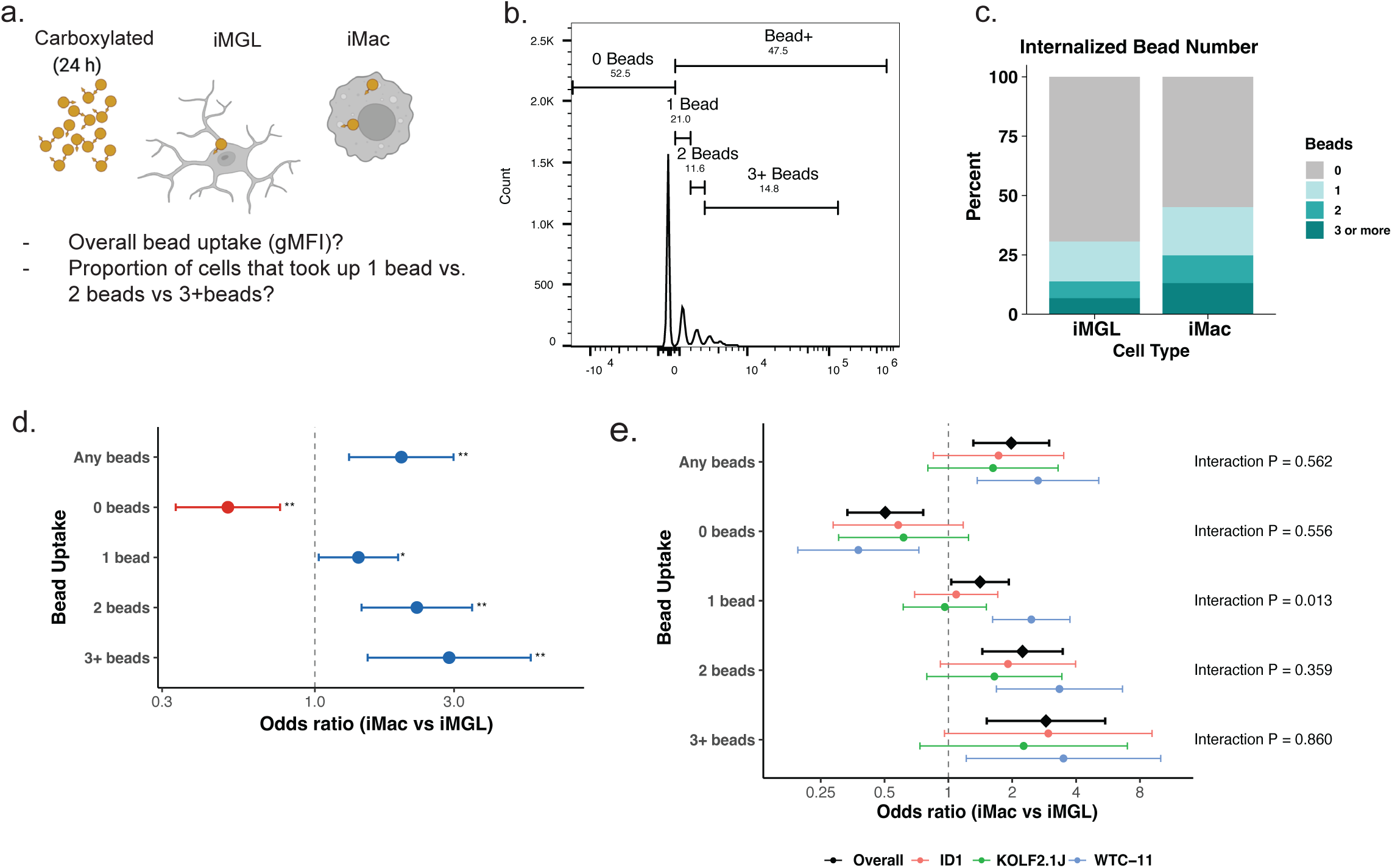
Quantitative analysis of bead uptake by flow cytometry. **(a)** Schematic overview of the carboxylated fluorescent bead uptake after 24 hours of treatment in iMGLs and iMacs. **(b)** Representative flow cytometry histogram showing discrete fluorescence intensity peaks used to gate and quantify cells containing 0, 1, 2, or 3+ beads, as well as total bead-positive (Bead+) cells. **(c)** Stacked bar graph illustrating the relative proportion of cells internalizing 0, 1, 2, or 3+ beads per cell in iMGLs compared to iMacs. **(d-e)** Forest plot showing odds ratios (iMac vs. iMGL) for overall bead uptake (Any beads) and individual bead counts (0, 1, 2, 3+ beads) **(d)** across all three donors and **(e)** stratified by iPSC donor line (ID1, KOLF2.1J, and WTC-11). Statistical significance and interaction effects were evaluated using generalized linear mixed-effects models. Full statistical details are provided in the Supplementary Statistical Report.

**Supplementary Figure 7.**
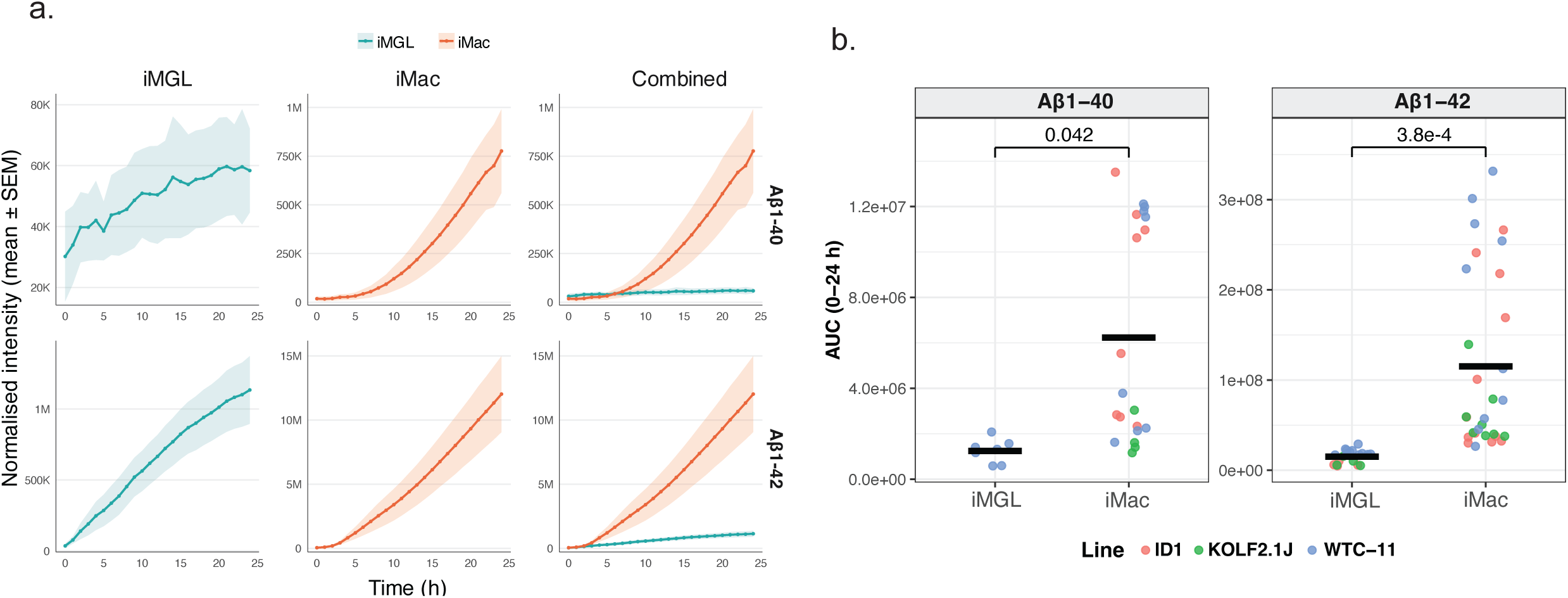
Differential phagocytic uptake of Aβ 1-40 and Aβ 1-42 between iMGLs and iMacs. **(a)** Time-course curves of normalized fluorescence intensity (mean ± SEM) showing uptake of Aβ 1-40 (top) and Aβ 1-42 (bottom) in iMGLs, iMacs, and overlaid comparisons (“Combined”) over 24 hours. Note that y-axes use different scales; uptake of Aβ 1-40 is minimal in iMGLs compared to iMacs, and uptake of Aβ 1-40 is minimal compared to Aβ 1-42. **(b)** Quantification of total amyloid uptake calculated as area under the curve (AUC) for Aβ 1-40 and Aβ 1-42. Statistical significance and interaction effects were evaluated using generalized linear mixed-effects models. Full statistical details are provided in the Supplementary Statistical Report.

