## Supplementary Statistical Reporting Document for "A rapid, genetically-tractable human iPSC-derived model for studying tissue-resident macrophage biology in Alzheimer’s and other diseases"

**Fig. 2A & E - Differential Expression Analysis of Bulk RNA-SEQ Data**

Differential expression was tested with a linear mixed model in dream (variancePartition) with cell type as a fixed effect and clone as a random effect (formula: ~ Cell_Type + (1 | Clone)), on 6 iMAC and 6 iMGL samples derived from three clones of the WTC11 iPSC line (WT-C, WT-D, WT-M; two samples per clone per cell type). Genes were tested individually; t-statistics are moderated (empirical Bayes) and P values were adjusted by Benjamini-Hochberg across all genes tested. Genes with P_adj < 0.05 and |log2FC| > 1 were considered differentially expressed (DEGs)

**Fig. 2B; Fig. 3B, F; Fig. 4C: Flow Cytometry Profiling of Surface Markers Between iMac and iMGL**

MFI values were rank-based inverse-normal transformed and scaled, then modelled with a linear mixed model (fixed: cell type, donor, differentiation; with a random intercept for culture - a combined “Donor_Cell_Type_Experiment” variable to account for technical replicates from the same donor line for each differentiation, with Kenward-Roger degrees of freedom. P values within each model are unadjusted; P_adj is Holm-adjusted across the 14 markers in this panel and is the value reported in the main text.

CD206 (Fig. 2B)

We fitted a linear mixed model (estimated using REML and nloptwrap optimizer) to predict

CD206_INT with Cell_Type, Donor and Differentiation (formula: CD206_INT ~ Cell_Type +

Donor + Differentiation). The model included Culture as random effect (formula: ~1 |

Culture). The model's total explanatory power is substantial (conditional R2 = 0.96) and

the part related to the fixed effects alone (marginal R2) is of 0.59. The model's

intercept, corresponding to Cell_Type = iMGL, Donor = ID1 and Differentiation = D1, is at

-0.82 (95% CI [-1.65, 0.02], t(12.00) = -2.12, p = 0.055). Within this model:

- The effect of Cell Type [iMac] is statistically significant and positive (beta = 1.59,

95% CI [0.91, 2.28], t(12.00) = 5.07, p < .001; Std. beta = 1.59, 95% CI [0.91, 2.28])

- The effect of Donor [KOLF] is statistically non-significant and positive (beta = 0.13,

95% CI [-0.71, 0.96], t(12.00) = 0.33, p = 0.746; Std. beta = 0.13, 95% CI [-0.71, 0.96])

- The effect of Donor [WTC11] is statistically non-significant and positive (beta = 0.30,

95% CI [-0.53, 1.14], t(12.00) = 0.79, p = 0.444; Std. beta = 0.30, 95% CI [-0.53, 1.14])

- The effect of Differentiation [D2] is statistically non-significant and negative (beta

= -0.12, 95% CI [-0.96, 0.72], t(12.00) = -0.32, p = 0.757; Std. beta = -0.12, 95% CI

[-0.96, 0.72])

- The effect of Differentiation [D3] is statistically non-significant and negative (beta

= -0.25, 95% CI [-1.09, 0.59], t(12.00) = -0.65, p = 0.528; Std. beta = -0.25, 95% CI

[-1.09, 0.59])

Parameters were scaled by the mean and the SD of the response variable. 95% Confidence

Intervals (CIs) and p-values were computed using a Wald t-distribution with Kenward-Roger

approximation.

Emmeans (inverse-normal-transformed, scaled MFI)

contrast	estimate	SE df lower.CL upper.CL t.ratio p.value

iMac - iMGL	1.59 0.314 12	0.908	2.28	5.073 0.0003

Results are averaged over the levels of: Donor, Differentiation

Degrees-of-freedom method: kenward-roger

Confidence level used: 0.95

Back-transformed to MFI (iMac - iMGL): β = 14,454, 95% CI: [8,877, 22,984]

Holm-adjusted across 14 markers: P_adj = 0.002

Model assumptions

OK: residuals appear as normally distributed (p = 0.943).

OK: Error variance appears to be homoscedastic (p = 0.953).

Singular fit (culture variance estimated at ~0): FALSE

\# Check for Multicollinearity

Low Correlation

Term VIF	VIF CI adj. VIF Tolerance

Cell_Type 1.00 [1.00, 1.00]	1.00	1.00

Donor 1.00 [1.00, 1.00]	1.00	1.00

Differentiation 1.00 [1.00, 1.00]	1.00	1.00

CD163 (Fig. 2B)

We fitted a linear mixed model (estimated using REML and nloptwrap optimizer) to predict

CD163_INT with Cell_Type, Donor and Differentiation (formula: CD163_INT ~ Cell_Type +

Donor + Differentiation). The model included Culture as random effect (formula: ~1 |

Culture). The model's total explanatory power is substantial (conditional R2 = 0.93) and

the part related to the fixed effects alone (marginal R2) is of 0.75. The model's

intercept, corresponding to Cell_Type = iMGL, Donor = ID1 and Differentiation = D1, is at

-0.29 (95% CI [-0.87, 0.30], t(12.00) = -1.07, p = 0.304). Within this model:

- The effect of Cell Type [iMac] is statistically significant and positive (beta = 1.59,

95% CI [1.11, 2.07], t(12.00) = 7.26, p < .001; Std. beta = 1.59, 95% CI [1.11, 2.07])

- The effect of Donor [KOLF] is statistically non-significant and negative (beta = -0.40,

95% CI [-0.99, 0.18], t(12.00) = -1.49, p = 0.161; Std. beta = -0.40, 95% CI [-0.99,

0.18])

- The effect of Donor [WTC11] is statistically non-significant and negative (beta =

-0.09, 95% CI [-0.67, 0.50], t(12.00) = -0.32, p = 0.751; Std. beta = -0.09, 95% CI

[-0.67, 0.50])

- The effect of Differentiation [D2] is statistically non-significant and negative (beta

= -0.21, 95% CI [-0.80, 0.37], t(12.00) = -0.78, p = 0.449; Std. beta = -0.21, 95% CI

[-0.80, 0.37])

- The effect of Differentiation [D3] is statistically significant and negative (beta =

-0.82, 95% CI [-1.41, -0.24], t(12.00) = -3.07, p = 0.010; Std. beta = -0.82, 95% CI

[-1.41, -0.24])

Parameters were scaled by the mean and the SD of the response variable. 95% Confidence

Intervals (CIs) and p-values were computed using a Wald t-distribution with Kenward-Roger

approximation.

Emmeans (inverse-normal-transformed, scaled MFI)

contrast	estimate	SE df lower.CL upper.CL t.ratio p.value

iMac - iMGL	1.59 0.219 12	1.11	2.07	7.260 <.0001

Results are averaged over the levels of: Donor, Differentiation

Degrees-of-freedom method: kenward-roger

Confidence level used: 0.95

Back-transformed to MFI (iMac - iMGL): β = 22,512, 95% CI: [18,201, 24,872]

Holm-adjusted across 14 markers: P_adj = 1.3 × 10^-4

Model assumptions

Warning: Non-normality of residuals detected (p < .001).

OK: Error variance appears to be homoscedastic (p = 0.849).

Singular fit (culture variance estimated at ~0): FALSE

\# Check for Multicollinearity

Low Correlation

Term VIF	VIF CI adj. VIF Tolerance

Cell_Type 1.00 [1.00, 1.00]	1.00	1.00

Donor 1.00 [1.00, 1.00]	1.00	1.00

Differentiation 1.00 [1.00, 1.00]	1.00	1.00

HLA.DR (Fig.2B)

We fitted a linear mixed model (estimated using REML and nloptwrap optimizer) to predict

HLA.DR_INT with Cell_Type, Donor and Differentiation (formula: HLA.DR_INT ~ Cell_Type +

Donor + Differentiation). The model included Culture as random effect (formula: ~1 |

Culture). The model's total explanatory power is substantial (conditional R2 = 0.98) and

the part related to the fixed effects alone (marginal R2) is of 0.41. The model's

intercept, corresponding to Cell_Type = iMGL, Donor = ID1 and Differentiation = D1, is at

0.60 (95% CI [-0.45, 1.66], t(12.00) = 1.24, p = 0.238). Within this model:

- The effect of Cell Type [iMac] is statistically significant and negative (beta = -1.39,

95% CI [-2.25, -0.53], t(12.00) = -3.51, p = 0.004; Std. beta = -1.39, 95% CI [-2.25,

-0.53])

- The effect of Donor [KOLF] is statistically non-significant and negative (beta = -0.03,

95% CI [-1.09, 1.02], t(12.00) = -0.07, p = 0.947; Std. beta = -0.03, 95% CI [-1.09,

1.02])

- The effect of Donor [WTC11] is statistically non-significant and negative (beta =

-0.03, 95% CI [-1.08, 1.03], t(12.00) = -0.05, p = 0.957; Std. beta = -0.03, 95% CI

[-1.08, 1.03])

- The effect of Differentiation [D2] is statistically non-significant and positive (beta

= 0.25, 95% CI [-0.80, 1.31], t(12.00) = 0.52, p = 0.610; Std. beta = 0.25, 95% CI

[-0.80, 1.31])

- The effect of Differentiation [D3] is statistically non-significant and positive (beta

= 0.08, 95% CI [-0.97, 1.14], t(12.00) = 0.18, p = 0.864; Std. beta = 0.08, 95% CI

[-0.97, 1.14])

Parameters were scaled by the mean and the SD of the response variable. 95% Confidence

Intervals (CIs) and p-values were computed using a Wald t-distribution with Kenward-Roger

approximation.

Emmeans (inverse-normal-transformed, scaled MFI)

contrast	estimate	SE df lower.CL upper.CL t.ratio p.value

iMac - iMGL	-1.39 0.395 12	-2.25	-0.527 -3.512 0.0043

Results are averaged over the levels of: Donor, Differentiation

Degrees-of-freedom method: kenward-roger

Confidence level used: 0.95

Back-transformed to MFI (iMac - iMGL): β = -10,988, 95% CI: [-15,277, -4,403]

Holm-adjusted across 15 markers: P_adj = 0.021

Model assumptions

OK: residuals appear as normally distributed (p = 0.514).

OK: Error variance appears to be homoscedastic (p = 0.971).

Singular fit (culture variance estimated at ~0): FALSE

\# Check for Multicollinearity

Low Correlation

Term VIF VIF 95% CI adj. VIF Tolerance Tolerance 95% CI

Cell_Type 1.00 [1.00, ]	1.00	1.00	[	, 1.00]

Donor 1.00 [1.00, Inf]	1.00	1.00	[0.00, 1.00]

Differentiation 1.00 [1.00, Inf]	1.00	1.00	[0.00, 1.00]

CD36 (Fig. 3B)

We fitted a linear mixed model (estimated using REML and nloptwrap optimizer) to predict

CD36_INT with Cell_Type, Donor and Differentiation (formula: CD36_INT ~ Cell_Type + Donor

+ Differentiation). The model included Culture as random effect (formula: ~1 | Culture).

The model's total explanatory power is substantial (conditional R2 = 0.95) and the part

related to the fixed effects alone (marginal R2) is of 0.71. The model's intercept,

corresponding to Cell_Type = iMGL, Donor = ID1 and Differentiation = D1, is at -0.49 (95%

CI [-1.15, 0.18], t(12.00) = -1.60, p = 0.135). Within this model:

- The effect of Cell Type [iMac] is statistically significant and positive (beta = 1.59,

95% CI [1.05, 2.13], t(12.00) = 6.41, p < .001; Std. beta = 1.59, 95% CI [1.05, 2.13])

- The effect of Donor [KOLF] is statistically non-significant and negative (beta = -0.46,

95% CI [-1.12, 0.21], t(12.00) = -1.50, p = 0.158; Std. beta = -0.46, 95% CI [-1.12,

0.21])

- The effect of Donor [WTC11] is statistically non-significant and positive (beta = 0.17,

95% CI [-0.50, 0.83], t(12.00) = 0.54, p = 0.598; Std. beta = 0.17, 95% CI [-0.50, 0.83])

- The effect of Differentiation [D2] is statistically non-significant and negative (beta

= -0.10, 95% CI [-0.76, 0.57], t(12.00) = -0.31, p = 0.760; Std. beta = -0.10, 95% CI

[-0.76, 0.57])

- The effect of Differentiation [D3] is statistically non-significant and negative (beta

= -0.54, 95% CI [-1.20, 0.13], t(12.00) = -1.76, p = 0.103; Std. beta = -0.54, 95% CI

[-1.20, 0.13])

Parameters were scaled by the mean and the SD of the response variable. 95% Confidence

Intervals (CIs) and p-values were computed using a Wald t-distribution with Kenward-Roger

approximation.

Emmeans (inverse-normal-transformed, scaled MFI)

contrast	estimate	SE df lower.CL upper.CL t.ratio p.value

iMac - iMGL	1.59 0.248 12	1.05	2.13	6.405 <.0001

Results are averaged over the levels of: Donor, Differentiation

Degrees-of-freedom method: kenward-roger

Confidence level used: 0.95

Back-transformed to MFI (iMac - iMGL): β = 77,495, 95% CI: [62,159, 85,403]

Holm-adjusted across 15 markers: P_adj = 3.7 × 10^-4

Model assumptions

Warning: Non-normality of residuals detected (p = 0.006).

OK: Error variance appears to be homoscedastic (p = 0.829).

Singular fit (culture variance estimated at ~0): FALSE

\# Check for Multicollinearity

Low Correlation

Term VIF	VIF CI adj. VIF Tolerance

Cell_Type 1.00 [1.00, 1.00]	1.00	1.00

Donor 1.00 [1.00, 1.00]	1.00	1.00

Differentiation 1.00 [1.00, 1.00]	1.00	1.00

CD64 (Fig. 3B)

We fitted a linear mixed model (estimated using REML and nloptwrap optimizer) to predict

CD64_INT with Cell_Type, Donor and Differentiation (formula: CD64_INT ~ Cell_Type + Donor

+ Differentiation). The model included Culture as random effect (formula: ~1 | Culture).

The model's total explanatory power is substantial (conditional R2 = 0.96) and the part

related to the fixed effects alone (marginal R2) is of 0.67. The model's intercept,

corresponding to Cell_Type = iMGL, Donor = ID1 and Differentiation = D1, is at -0.23 (95%

CI [-0.95, 0.49], t(12.00) = -0.70, p = 0.497). Within this model:

- The effect of Cell Type [iMac] is statistically significant and positive (beta = 1.59,

95% CI [1.00, 2.18], t(12.00) = 5.90, p < .001; Std. beta = 1.59, 95% CI [1.00, 2.18])

- The effect of Donor [KOLF] is statistically non-significant and negative (beta = -0.58,

95% CI [-1.30, 0.14], t(12.00) = -1.76, p = 0.104; Std. beta = -0.58, 95% CI [-1.30,

0.14])

- The effect of Donor [WTC11] is statistically non-significant and negative (beta =

-0.29, 95% CI [-1.01, 0.43], t(12.00) = -0.89, p = 0.392; Std. beta = -0.29, 95% CI

[-1.01, 0.43])

- The effect of Differentiation [D2] is statistically non-significant and negative (beta

= -0.49, 95% CI [-1.21, 0.23], t(12.00) = -1.49, p = 0.162; Std. beta = -0.49, 95% CI

[-1.21, 0.23])

- The effect of Differentiation [D3] is statistically non-significant and negative (beta

= -0.33, 95% CI [-1.05, 0.39], t(12.00) = -0.99, p = 0.341; Std. beta = -0.33, 95% CI

[-1.05, 0.39])

Parameters were scaled by the mean and the SD of the response variable. 95% Confidence

Intervals (CIs) and p-values were computed using a Wald t-distribution with Kenward-Roger

approximation.

Emmeans (inverse-normal-transformed, scaled MFI)

contrast	estimate	SE df lower.CL upper.CL t.ratio p.value

iMac - iMGL	1.59 0.27 12	1	2.18	5.900 0.0001

Results are averaged over the levels of: Donor, Differentiation

Degrees-of-freedom method: kenward-roger

Confidence level used: 0.95

Back-transformed to MFI (iMac - iMGL): β = 40,247, 95% CI: [33,811, 53,052]

Holm-adjusted across 14 markers: P_adj = 7.3 × 10^-4

Model assumptions

Warning: Non-normality of residuals detected (p = 0.007).

OK: Error variance appears to be homoscedastic (p = 0.992).

Singular fit (culture variance estimated at ~0): FALSE

\# Check for Multicollinearity

Low Correlation

Term VIF	VIF CI adj. VIF Tolerance

Cell_Type 1.00 [1.00, 1.00]	1.00	1.00

Donor 1.00 [1.00, 1.00]	1.00	1.00

Differentiation 1.00 [1.00, 1.00]	1.00	1.00

CLEC7A (Fig. 3B)

We fitted a linear mixed model (estimated using REML and nloptwrap optimizer) to predict

CLEC7A_INT with Cell_Type, Donor and Differentiation (formula: CLEC7A_INT ~ Cell_Type +

Donor + Differentiation). The model included Culture as random effect (formula: ~1 |

Culture). The model's total explanatory power is substantial (conditional R2 = 0.89) and

the part related to the fixed effects alone (marginal R2) is of 0.77. The model's

intercept, corresponding to Cell_Type = iMGL, Donor = ID1 and Differentiation = D1, is at

-0.20 (95% CI [-0.70, 0.31], t(12.00) = -0.86, p = 0.407). Within this model:

- The effect of Cell Type [iMac] is statistically significant and positive (beta = 1.59,

95% CI [1.18, 2.00], t(12.00) = 8.40, p < .001; Std. beta = 1.59, 95% CI [1.18, 2.00])

- The effect of Donor [KOLF] is statistically non-significant and negative (beta = -0.21,

95% CI [-0.71, 0.30], t(12.00) = -0.89, p = 0.392; Std. beta = -0.21, 95% CI [-0.71,

0.30])

- The effect of Donor [WTC11] is statistically significant and negative (beta = -0.85,

95% CI [-1.35, -0.34], t(12.00) = -3.65, p = 0.003; Std. beta = -0.85, 95% CI [-1.35,

-0.34])

- The effect of Differentiation [D2] is statistically non-significant and negative (beta

= -0.31, 95% CI [-0.82, 0.19], t(12.00) = -1.36, p = 0.200; Std. beta = -0.31, 95% CI

[-0.82, 0.19])

- The effect of Differentiation [D3] is statistically non-significant and negative (beta

= -0.42, 95% CI [-0.93, 0.08], t(12.00) = -1.82, p = 0.094; Std. beta = -0.42, 95% CI

[-0.93, 0.08])

Parameters were scaled by the mean and the SD of the response variable. 95% Confidence

Intervals (CIs) and p-values were computed using a Wald t-distribution with Kenward-Roger

approximation.

Emmeans (inverse-normal-transformed, scaled MFI)

contrast	estimate	SE df lower.CL upper.CL t.ratio p.value

iMac - iMGL	1.59 0.189 12	1.18	2	8.404 <.0001

Results are averaged over the levels of: Donor, Differentiation

Degrees-of-freedom method: kenward-roger

Confidence level used: 0.95

Back-transformed to MFI (iMac - iMGL): β = 5,089, 95% CI: [4,445, 6,141]

Holm-adjusted across 15 markers: P_adj = 3.2 × 10^-5

Model assumptions

Warning: Non-normality of residuals detected (p < .001).

OK: Error variance appears to be homoscedastic (p = 0.795).

Singular fit (culture variance estimated at ~0): FALSE

\# Check for Multicollinearity

Low Correlation

Term VIF	VIF CI adj. VIF Tolerance

Cell_Type 1.00 [1.00, 1.00]	1.00	1.00

Donor 1.00 [1.00, 1.00]	1.00	1.00

Differentiation 1.00 [1.00, 1.00]	1.00	1.00

LAMP1 (Fig. 3F)

We fitted a linear mixed model (estimated using REML and nloptwrap optimizer) to predict

LAMP1_INT with Cell_Type, Donor and Differentiation (formula: LAMP1_INT ~ Cell_Type +

Donor + Differentiation). The model included Culture as random effect (formula: ~1 |

Culture). The model's total explanatory power is substantial (conditional R2 = 0.91) and

the part related to the fixed effects alone (marginal R2) is of 0.63. The model's

intercept, corresponding to Cell_Type = iMGL, Donor = ID1 and Differentiation = D1, is at

-0.86 (95% CI [-1.60, -0.13], t(12.00) = -2.57, p = 0.025). Within this model:

- The effect of Cell Type [iMac] is statistically significant and positive (beta = 1.60,

95% CI [1.00, 2.19], t(12.00) = 5.82, p < .001; Std. beta = 1.60, 95% CI [1.00, 2.19])

- The effect of Donor [KOLF] is statistically non-significant and negative (beta = -0.18,

95% CI [-0.91, 0.55], t(12.00) = -0.53, p = 0.603; Std. beta = -0.18, 95% CI [-0.91,

0.55])

- The effect of Donor [WTC11] is statistically non-significant and positive (beta = 0.27,

95% CI [-0.46, 1.01], t(12.00) = 0.82, p = 0.430; Std. beta = 0.27, 95% CI [-0.46, 1.01])

- The effect of Differentiation [D2] is statistically non-significant and positive (beta

= 0.21, 95% CI [-0.52, 0.94], t(12.00) = 0.62, p = 0.546; Std. beta = 0.21, 95% CI

[-0.52, 0.94])

- The effect of Differentiation [D3] is statistically non-significant and negative (beta

= -0.11, 95% CI [-0.84, 0.62], t(12.00) = -0.32, p = 0.754; Std. beta = -0.11, 95% CI

[-0.84, 0.62])

Parameters were scaled by the mean and the SD of the response variable. 95% Confidence

Intervals (CIs) and p-values were computed using a Wald t-distribution with Kenward-Roger

approximation.

Emmeans (inverse-normal-transformed, scaled MFI)

contrast	estimate	SE df lower.CL upper.CL t.ratio p.value

iMac - iMGL	1.6 0.274 12	0.998	2.19	5.818 0.0001

Results are averaged over the levels of: Donor, Differentiation

Degrees-of-freedom method: kenward-roger

Confidence level used: 0.95

Back-transformed to MFI (iMac - iMGL): β = 294,168, 95% CI: [242,778, 327,936]

Holm-adjusted across 14 markers: P_adj = 7.3 × 10^-4

Model assumptions

OK: residuals appear as normally distributed (p = 0.114).

OK: Error variance appears to be homoscedastic (p = 0.978).

Singular fit (culture variance estimated at ~0): FALSE

\# Check for Multicollinearity

Low Correlation

Term VIF	VIF CI adj. VIF Tolerance

Cell_Type 1.00 [1.00, 1.00]	1.00	1.00

Donor 1.00 [1.00, 1.00]	1.00	1.00

Differentiation 1.00 [1.00, 1.00]	1.00	1.00

CD68 (Fig. 3F)

We fitted a linear mixed model (estimated using REML and nloptwrap optimizer) to predict

CD68_INT with Cell_Type, Donor and Differentiation (formula: CD68_INT ~ Cell_Type + Donor

+ Differentiation). The model included Culture as random effect (formula: ~1 | Culture).

The model's total explanatory power is substantial (conditional R2 = 0.97) and the part

related to the fixed effects alone (marginal R2) is of 0.37. The model's intercept,

corresponding to Cell_Type = iMGL, Donor = ID1 and Differentiation = D1, is at 0.47 (95%

CI [-0.63, 1.57], t(12.00) = 0.94, p = 0.367). Within this model:

- The effect of Cell Type [iMac] is statistically non-significant and positive (beta =

0.53, 95% CI [-0.37, 1.42], t(12.00) = 1.28, p = 0.224; Std. beta = 0.53, 95% CI [-0.37,

1.42])

- The effect of Donor [KOLF] is statistically non-significant and negative (beta = -0.73,

95% CI [-1.83, 0.36], t(12.00) = -1.45, p = 0.172; Std. beta = -0.73, 95% CI [-1.83,

0.36])

- The effect of Donor [WTC11] is statistically non-significant and negative (beta =

-0.33, 95% CI [-1.42, 0.77], t(12.00) = -0.65, p = 0.531; Std. beta = -0.33, 95% CI

[-1.42, 0.77])

- The effect of Differentiation [D2] is statistically non-significant and negative (beta

= -7.61e-03, 95% CI [-1.10, 1.09], t(12.00) = -0.02, p = 0.988; Std. beta = -7.61e-03,

95% CI [-1.10, 1.09])

- The effect of Differentiation [D3] is statistically significant and negative (beta =

-1.14, 95% CI [-2.24, -0.04], t(12.00) = -2.27, p = 0.043; Std. beta = -1.14, 95% CI

[-2.24, -0.04])

Parameters were scaled by the mean and the SD of the response variable. 95% Confidence

Intervals (CIs) and p-values were computed using a Wald t-distribution with Kenward-Roger

approximation.

Emmeans (inverse-normal-transformed, scaled MFI)

contrast	estimate	SE df lower.CL upper.CL t.ratio p.value

iMac - iMGL	0.527 0.411 12	-0.369	1.42	1.281 0.2243

Results are averaged over the levels of: Donor, Differentiation

Degrees-of-freedom method: kenward-roger

Confidence level used: 0.95

Back-transformed to MFI (iMac - iMGL): β = 16,693, 95% CI: [-8,865, 76,074]

Holm-adjusted across 14 markers: P_adj = 0.442

Model assumptions

OK: residuals appear as normally distributed (p = 0.187).

OK: Error variance appears to be homoscedastic (p = 0.937).

Singular fit (culture variance estimated at ~0): FALSE

\# Check for Multicollinearity

Low Correlation

Term VIF	VIF CI adj. VIF Tolerance

Cell_Type 1.00 [1.00, 1.00]	1.00	1.00

Donor 1.00 [1.00, 1.00]	1.00	1.00

Differentiation 1.00 [1.00, 1.00]	1.00	1.00

TREM2 (Fig. 4C)

We fitted a linear mixed model (estimated using REML and nloptwrap optimizer) to predict

TREM2_INT with Cell_Type, Donor and Differentiation (formula: TREM2_INT ~ Cell_Type +

Donor + Differentiation). The model included Culture as random effect (formula: ~1 |

Culture). The model's total explanatory power is substantial (conditional R2 = 0.97) and

the part related to the fixed effects alone (marginal R2) is of 0.65. The model's

intercept, corresponding to Cell_Type = iMGL, Donor = ID1 and Differentiation = D1, is at

-0.33 (95% CI [-1.08, 0.43], t(12.00) = -0.94, p = 0.364). Within this model:

- The effect of Cell Type [iMac] is statistically significant and positive (beta = 1.59,

95% CI [0.97, 2.21], t(12.00) = 5.62, p < .001; Std. beta = 1.59, 95% CI [0.97, 2.21])

- The effect of Donor [KOLF] is statistically non-significant and negative (beta = -0.32,

95% CI [-1.07, 0.44], t(12.00) = -0.91, p = 0.380; Std. beta = -0.32, 95% CI [-1.07,

0.44])

- The effect of Donor [WTC11] is statistically non-significant and negative (beta =

-0.05, 95% CI [-0.80, 0.71], t(12.00) = -0.13, p = 0.897; Std. beta = -0.05, 95% CI

[-0.80, 0.71])

- The effect of Differentiation [D2] is statistically non-significant and negative (beta

= -0.59, 95% CI [-1.35, 0.16], t(12.00) = -1.71, p = 0.113; Std. beta = -0.59, 95% CI

[-1.35, 0.16])

- The effect of Differentiation [D3] is statistically non-significant and negative (beta

= -0.45, 95% CI [-1.21, 0.30], t(12.00) = -1.30, p = 0.217; Std. beta = -0.45, 95% CI

[-1.21, 0.30])

Parameters were scaled by the mean and the SD of the response variable. 95% Confidence

Intervals (CIs) and p-values were computed using a Wald t-distribution with Kenward-Roger

approximation.

Emmeans (inverse-normal-transformed, scaled MFI)

contrast	estimate	SE df lower.CL upper.CL t.ratio p.value

iMac - iMGL	1.59 0.283 12	0.975	2.21	5.623 0.0001

Results are averaged over the levels of: Donor, Differentiation

Degrees-of-freedom method: kenward-roger

Confidence level used: 0.95

Back-transformed to MFI (iMac - iMGL): β = 12,347, 95% CI: [8,207, 13,846]

Holm-adjusted across 15 markers: P_adj = 7.8 × 10^-4

Model assumptions

OK: residuals appear as normally distributed (p = 0.318).

OK: Error variance appears to be homoscedastic (p = 0.965).

Singular fit (culture variance estimated at ~0): FALSE

\# Check for Multicollinearity

Low Correlation

Term VIF VIF 95% CI adj. VIF Tolerance Tolerance 95% CI

Cell_Type 1.00 [1.00, ]	1.00	1.00	[	, 1.00]

Donor 1.00 [1.00, Inf]	1.00	1.00	[0.00, 1.00]

Differentiation 1.00 [1.00, Inf]	1.00	1.00	[0.00, 1.00]

**Fig. 2C, D; Fig. 3A; Supp. Fig 4 - Gene Set Enrichment Analysis**

Gene-set enrichment analysis against databases was performed with fgsea (multilevel method) on all genes tested in the differential expression analysis (Figure 3A; 6 iMAC vs 6 iMGL, three WTC11 clones), ranked by the dream moderated t-statistic for iMAC vs iMGL. Gene sets were taken from MSigDB (v7.5.1) collections C2 (chemical and genetic perturbations; canonical pathways: Reactome, KEGG, WikiPathways, BioCarta, PID) and C5 (GO biological process, cellular component and molecular function; Human Phenotype Ontology). 23,478 gene sets of 1-1,725 genes (after matching to the ranked list) were tested, and P values were adjusted by Benjamini-Hochberg across all 23,478 sets jointly. Pathways with P_adj < 0.10 were considered significant. Published microglial and macrophage state signatures, including disease-associated microglia (compiled in Podlesny-Drabiniok et al., 2026), were tested separately with clusterProfiler GSEA on the same ranked list; [N] signatures were tested, with Benjamini-Hochberg adjustment within this collection only and significance at P_adj < [0.10]. Positive normalised enrichment score (NES) = enriched among genes higher in iMAC for both analyses.

**Figure 2E: iMAC vs iMGL - risk gene overlap with differentially expressed genes**

Differentially expressed genes (DEGs) were defined as FDR < 0.05 and |log2FC| > 1 in the

iMAC vs iMGL comparison, up- and downregulated combined (k = 4,136 DEGs: 2,678 upregulated and 1,458 downregulated in iMAC). The background (universe) was all genes tested in the differential expression analysis (N = 14,587); risk genes absent from this set were

excluded from each list before testing. Over-representation was tested with a one-sided

hypergeometric test, P(X >= observed overlap), which is identical to a one-sided Fisher's

exact test on the same 2x2 table. Odds ratios are conditional maximum-likelihood estimates

with two-sided exact 95% confidence intervals. Fold enrichment is the proportion of the

list that is differentially expressed divided by the background DEG rate (4,136/14,587 =

28.4%). P values are unadjusted; P_adj is Benjamini-Hochberg adjusted across the four risk

gene sets and is the value reported in the main text.

**Fig 3C; Supplementary Fig. 6D-E - Phagocytosis Uptake of Latex Beads (gMFI)**

gMFI

gMFI values were analysed untransformed unless residual diagnostics indicated that model assumptions were not met (transformation used: AllCells_gMFI = identity; BeadPositive_gMFI = untransformed); values were scaled, then modelled with a linear mixed model (fixed: cell type, donor, experiment; random intercept: culture, grouping each culture's replicate wells), with Kenward-Roger degrees of freedom. P values within each model are unadjusted; P_adj is Holm-adjusted across the 2 gMFI readouts and is the value reported in the main text.

Excluded wells (technical failure): WTC-11 iMGL 3 | 1.1

We fitted a linear mixed model (estimated using REML and nloptwrap optimizer)

to predict AllCells_identity with Cell_Type, Donor and Experiment (formula:

AllCells_identity ~ Cell_Type + Donor + Experiment). The model included Culture

as random effect (formula: ~1 | Culture). The model's total explanatory power

is substantial (conditional R2 = 0.96) and the part related to the fixed

effects alone (marginal R2) is of 0.56. The model's intercept, corresponding to

Cell_Type = iMGL, Donor = ID1 and Experiment = 1.1, is at 0.41 (95% CI [-0.46,

1.28], t(13.94) = 1.02, p = 0.324). Within this model:

- The effect of Cell Type [iMac] is statistically significant and positive

(beta = 0.75, 95% CI [0.08, 1.42], t(14.03) = 2.39, p = 0.032; Std. beta =

0.75, 95% CI [0.08, 1.42])

- The effect of Donor [KOLF] is statistically non-significant and positive

(beta = 0.32, 95% CI [-0.52, 1.15], t(13.90) = 0.82, p = 0.428; Std. beta =

0.32, 95% CI [-0.52, 1.15])

- The effect of Donor [WTC-11] is statistically non-significant and negative

(beta = -0.57, 95% CI [-1.43, 0.28], t(14.09) = -1.44, p = 0.171; Std. beta =

-0.57, 95% CI [-1.43, 0.28])

- The effect of Experiment [2.1] is statistically non-significant and negative

(beta = -0.71, 95% CI [-1.64, 0.22], t(14.00) = -1.64, p = 0.123; Std. beta =

-0.71, 95% CI [-1.64, 0.22])

- The effect of Experiment [2.2] is statistically non-significant and negative

(beta = -0.82, 95% CI [-1.94, 0.31], t(14.12) = -1.56, p = 0.142; Std. beta =

-0.82, 95% CI [-1.94, 0.31])

- The effect of Experiment [3.1] is statistically non-significant and negative

(beta = -0.89, 95% CI [-1.82, 0.04], t(14.03) = -2.06, p = 0.058; Std. beta =

-0.89, 95% CI [-1.82, 0.04])

- The effect of Experiment [4.1] is statistically significant and negative

(beta = -1.80, 95% CI [-2.93, -0.68], t(14.12) = -3.44, p = 0.004; Std. beta =

-1.80, 95% CI [-2.93, -0.68])

Parameters were scaled by the mean and the SD of the response variable. 95%

Confidence Intervals (CIs) and p-values were computed using a Wald

t-distribution with Kenward-Roger approximation.

Emmeans (identity-transformed, scaled gMFI)

contrast	estimate	SE df lower.CL upper.CL t.ratio p.value

iMac - iMGL	0.748 0.314 14	0.076	1.42	2.387 0.0316

Results are averaged over the levels of: Donor, Experiment

Degrees-of-freedom method: kenward-roger

Confidence level used: 0.95

Back-transformed to gMFI (iMac - iMGL): β = 346, 95% CI: [56, 635]; fold change = 1.52 [1.07, 2.36]

Holm-adjusted across 2 gMFI readouts: P_adj = 0.032

Model assumptions

OK: residuals appear as normally distributed (p = 0.075).

OK: Error variance appears to be homoscedastic (p = 0.968).

Singular fit (culture variance estimated at ~0): FALSE

\# Check for Multicollinearity

Low Correlation

Term VIF	VIF 95% CI adj. VIF Tolerance Tolerance 95% CI

Cell_Type 1.07 [1.00, 3.35]	1.03	0.94	[0.30, 1.00]

Donor 1.28 [1.09, 1.89]	1.13	0.78	[0.53, 0.92]

Experiment 1.33 [1.12, 1.93]	1.15	0.75	[0.52, 0.90]

BeadPositive

We fitted a linear mixed model (estimated using REML and nloptwrap optimizer)

to predict BeadPositive_identity with Cell_Type, Donor and Experiment (formula:

BeadPositive_identity ~ Cell_Type + Donor + Experiment). The model included

Culture as random effect (formula: ~1 | Culture). The model's total explanatory

power is substantial (conditional R2 = 0.96) and the part related to the fixed

effects alone (marginal R2) is of 0.61. The model's intercept, corresponding to

Cell_Type = iMGL, Donor = ID1 and Experiment = 1.1, is at 0.41 (95% CI [-0.39,

1.22], t(13.94) = 1.10, p = 0.289). Within this model:

- The effect of Cell Type [iMac] is statistically significant and positive

(beta = 0.86, 95% CI [0.24, 1.49], t(14.03) = 2.98, p = 0.010; Std. beta =

0.86, 95% CI [0.24, 1.49])

- The effect of Donor [KOLF] is statistically non-significant and positive

(beta = 0.36, 95% CI [-0.41, 1.13], t(13.89) = 1.01, p = 0.332; Std. beta =

0.36, 95% CI [-0.41, 1.13])

- The effect of Donor [WTC-11] is statistically non-significant and negative

(beta = -0.58, 95% CI [-1.37, 0.21], t(14.10) = -1.57, p = 0.139; Std. beta =

-0.58, 95% CI [-1.37, 0.21])

- The effect of Experiment [2.1] is statistically significant and negative

(beta = -0.89, 95% CI [-1.75, -0.03], t(14.00) = -2.21, p = 0.044; Std. beta =

-0.89, 95% CI [-1.75, -0.03])

- The effect of Experiment [2.2] is statistically non-significant and negative

(beta = -0.95, 95% CI [-1.99, 0.09], t(14.13) = -1.95, p = 0.071; Std. beta =

-0.95, 95% CI [-1.99, 0.09])

- The effect of Experiment [3.1] is statistically significant and negative

(beta = -0.99, 95% CI [-1.85, -0.14], t(14.03) = -2.48, p = 0.026; Std. beta =

-0.99, 95% CI [-1.85, -0.14])

- The effect of Experiment [4.1] is statistically significant and negative

(beta = -1.70, 95% CI [-2.74, -0.66], t(14.13) = -3.50, p = 0.003; Std. beta =

-1.70, 95% CI [-2.74, -0.66])

Parameters were scaled by the mean and the SD of the response variable. 95%

Confidence Intervals (CIs) and p-values were computed using a Wald

t-distribution with Kenward-Roger approximation.

Emmeans (identity-transformed, scaled gMFI)

contrast	estimate	SE df lower.CL upper.CL t.ratio p.value

iMac - iMGL	0.864 0.29 14	0.242	1.49	2.978 0.0100

Results are averaged over the levels of: Donor, Experiment

Degrees-of-freedom method: kenward-roger

Confidence level used: 0.95

Back-transformed to gMFI (iMac - iMGL): β = 381, 95% CI: [127, 637]; fold change = 1.18 [1.06, 1.33]

Holm-adjusted across 2 gMFI readouts: P_adj = 0.020

Model assumptions

Warning: Non-normality of residuals detected (p = 0.025).

OK: Error variance appears to be homoscedastic (p = 0.982).

Singular fit (culture variance estimated at ~0): FALSE

\# Check for Multicollinearity

Low Correlation

Term VIF	VIF 95% CI adj. VIF Tolerance Tolerance 95% CI

Cell_Type 1.07 [1.00, 3.34]	1.03	0.94	[0.30, 1.00]

Donor 1.28 [1.09, 1.89]	1.13	0.78	[0.53, 0.92]

Experiment 1.33 [1.12, 1.93]	1.15	0.75	[0.52, 0.90]

**Figure 3D; Supplementary Fig.7B - Amyloid Phagocytosis**

Time Course - AB42 and AB40

For each substrate, normalised intensity was modelled with a generalised linear mixed model (glmmTMB; family: AB-40 = tweedie; AB-42 = tweedie) with cell type, a natural spline of time (3 df) and their interaction, line and experiment as fixed effects; random intercepts for culture (grouping each culture's replicate wells) and a random intercept and time slope per well (repeated measures). Families were chosen from simulated-residual (DHARMa) diagnostics across compatible families; no transformation was applied. Effects were tested by likelihood-ratio tests; iMac - iMGL differences at 4, 8, 12, 16, 20, 24 h are Holm-adjusted across timepoints.

Excluded wells: none

Family comparison

|Substrate |family |converged | AIC| Uniformity_p| Dispersion_p| Outliers_p| ZeroInflation_p|

|AB-40 |tweedie |TRUE | 14446.91| 0| 0.880| 0.38| NA|

|AB-40 |gamma |TRUE | 14984.13| 0| 0.704| 0.08| NA|

|AB-40 |gaussian_disp |FALSE | NA| 0| 0.000| 0.14| NA|

|AB-42 |tweedie |TRUE | 33783.49| 0| 0.720| 0.56| 0.552|

|AB-42 |hurdle_gamma |TRUE | 34691.83| 0| 0.000| 0.04| 1.000|

|AB-42 |gaussian_disp |FALSE | NA| 0| 0.000| 0.00| 0.000|

AB-40

Time (main + interaction): chi2(6) = 954.30, P = 6.8 × 10^-203

Cell type x time interaction: chi2(3) = 222.82, P = 4.9 × 10^-48

Overall cell type effect (main + interaction): chi2(4) = 263.23, P = 9.2 × 10^-56

iMac - iMGL difference in normalised intensity by timepoint (Holm across timepoints)

| Elapsed| estimate| SE| asymp.LCL| asymp.UCL| p.value| p_adj|Significance |

| 4| -20886.375| 15530.36| -51325.33| 9552.58| 0.179| 0.357|ns |

| 8| -9053.418| 16703.29| -41791.26| 23684.42| 0.588| 0.588|ns |

| 12| 41835.819| 21829.01| -948.25| 84619.89| 0.055| 0.166|ns |

| 16| 171339.078| 35924.29| 100928.76| 241749.40| 0.000| 0.000|*** |

| 20| 361687.053| 46280.37| 270979.19| 452394.91| 0.000| 0.000|*** |

| 24| 587531.082| 44596.49| 500123.57| 674938.59| 0.000| 0.000|*** |

Fixed effects (model scale)

Estimate Std. Error z value Pr(>|z|)

(Intercept)	10.0172	0.622 16.118 1.92e-58

Cell_TypeiMac	-2.1769	0.713 -3.053 2.26e-03

ns(Elapsed, 3)1	0.4556	0.302	1.509 1.31e-01

ns(Elapsed, 3)2	1.0128	0.617	1.640 1.01e-01

ns(Elapsed, 3)3	0.4218	0.424	0.995 3.20e-01

LineKOLF	-0.4815	0.130 -3.718 2.01e-04

LineWTC-11	0.0856	0.101	0.844 3.98e-01

Exp4.1	0.3542	0.148	2.386 1.70e-02

Cell_TypeiMac:ns(Elapsed, 3)1	3.5620	0.351 10.141 3.65e-24

Cell_TypeiMac:ns(Elapsed, 3)2	6.0742	0.719	8.450 2.92e-17

Cell_TypeiMac:ns(Elapsed, 3)3	3.7921	0.493	7.696 1.40e-14

Model diagnostics (DHARMa): uniformity P = 6.1 × 10^-10; dispersion P = 0.880; outliers P = 0.380

AB-42

Time (main + interaction): chi2(6) = 1555.21, P = 0.0e+00

Cell type x time interaction: chi2(3) = 102.92, P = 3.7 × 10^-22

Overall cell type effect (main + interaction): chi2(4) = 132.02, P = 1.4 × 10^-27

iMac - iMGL difference in normalised intensity by timepoint (Holm across timepoints)

| Elapsed| estimate| SE| asymp.LCL| asymp.UCL| p.value| p_adj|Significance |

| 4| 40876.31| 57127.12| -71090.8| 152843.4| 0.474| 0.474|ns |

| 8| 587516.36| 222373.61| 151672.1| 1023360.6| 0.008| 0.016|* |

| 12| 1986174.01| 488854.05| 1028037.7| 2944310.3| 0.000| 0.000|*** |

| 16| 3782636.80| 709257.84| 2392517.0| 5172756.6| 0.000| 0.000|*** |

| 20| 6278646.87| 978463.38| 4360893.9| 8196399.8| 0.000| 0.000|*** |

| 24| 10069572.08| 1502737.26| 7124261.2| 13014883.0| 0.000| 0.000|*** |

Fixed effects (model scale)

Estimate Std. Error z value Pr(>|z|)

(Intercept)	10.018	0.446 22.443 1.50e-111

Cell_TypeiMac	-0.601	0.521 -1.155 2.48e-01

ns(Elapsed, 3)1	2.338	0.231 10.133 3.94e-24

ns(Elapsed, 3)2	5.577	0.487 11.457 2.18e-30

ns(Elapsed, 3)3	1.922	0.314	6.110 9.94e-10

LineKOLF	0.105	0.288	0.365 7.15e-01

LineWTC-11	0.574	0.251	2.289 2.21e-02

Exp3.1	-0.176	0.255 -0.689 4.91e-01

Exp4.1	0.476	0.283	1.682 9.26e-02

Cell_TypeiMac:ns(Elapsed, 3)1	1.978	0.293	6.758 1.40e-11

Cell_TypeiMac:ns(Elapsed, 3)2	4.447	0.617	7.205 5.80e-13

Cell_TypeiMac:ns(Elapsed, 3)3	1.990	0.401	4.961 7.02e-07

Model diagnostics (DHARMa): uniformity P = 5.4 × 10^-5; dispersion P = 0.720; outliers P = 0.560

AUC

Per-well area under the uptake curve (0-24 h, trapezoidal) was analysed untransformed unless residual diagnostics indicated that model assumptions were not met (transformation used: AB-40 = log; AB-42 = log), scaled, and modelled per substrate with a linear mixed model (fixed: cell type, line, experiment; random intercept: culture, grouping each culture's replicate wells), with Kenward-Roger degrees of freedom. P values within each model are unadjusted; P_adj is Holm-adjusted across the two amyloid substrates.

Excluded wells: none

Design (wells per substrate, cell type, line and experiment)

|Substrate |Cell_Type |Line |Exp | n|

|AB-40 |iMGL |WTC-11 |3.1 | 3|

|AB-40 |iMGL |WTC-11 |4.1 | 4|

|AB-40 |iMac |ID1 |3.1 | 4|

|AB-40 |iMac |ID1 |4.1 | 4|

|AB-40 |iMac |KOLF |3.1 | 4|

|AB-40 |iMac |WTC-11 |3.1 | 4|

|AB-40 |iMac |WTC-11 |4.1 | 4|

|AB-42 |iMGL |ID1 |1.1 | 4|

|AB-42 |iMGL |KOLF |1.1 | 4|

|AB-42 |iMGL |WTC-11 |1.1 | 4|

|AB-42 |iMGL |WTC-11 |3.1 | 3|

|AB-42 |iMGL |WTC-11 |4.1 | 4|

|AB-42 |iMac |ID1 |1.1 | 4|

|AB-42 |iMac |ID1 |3.1 | 4|

|AB-42 |iMac |ID1 |4.1 | 4|

|AB-42 |iMac |KOLF |1.1 | 4|

|AB-42 |iMac |KOLF |3.1 | 4|

|AB-42 |iMac |WTC-11 |1.1 | 2|

|AB-42 |iMac |WTC-11 |3.1 | 4|

|AB-42 |iMac |WTC-11 |4.1 | 4|

AB-40

We fitted a linear mixed model (estimated using REML and nloptwrap optimizer) to predict

AB40_AUC_log with Cell_Type, Line and Exp (formula: AB40_AUC_log ~ Cell_Type + Line +

Exp). The model included Culture as random effect (formula: ~1 | Culture). The model's

total explanatory power is substantial (conditional R2 = 0.91) and the part related to

the fixed effects alone (marginal R2) is of 0.82. The model's intercept, corresponding to

Cell_Type = iMGL, Line = ID1 and Exp = 3.1, is at -1.53 (95% CI [-3.40, 0.34], t(2.03) =

-3.49, p = 0.072). Within this model:

- The effect of Cell Type [iMac] is statistically significant and positive (beta = 1.59,

95% CI [0.15, 3.04], t(2.02) = 4.69, p = 0.042; Std. beta = 1.59, 95% CI [0.15, 3.04])

- The effect of Line [KOLF] is statistically non-significant and negative (beta = -0.68,

95% CI [-2.59, 1.23], t(1.96) = -1.57, p = 0.261; Std. beta = -0.68, 95% CI [-2.59,

1.23])

- The effect of Line [WTC-11] is statistically non-significant and negative (beta =

-0.14, 95% CI [-1.63, 1.34], t(1.95) = -0.43, p = 0.711; Std. beta = -0.14, 95% CI

[-1.63, 1.34])

- The effect of Exp [4.1] is statistically significant and positive (beta = 1.21, 95% CI

[0.02, 2.41], t(2.00) = 4.38, p = 0.048; Std. beta = 1.21, 95% CI [0.02, 2.41])

Parameters were scaled by the mean and the SD of the response variable. 95% Confidence

Intervals (CIs) and p-values were computed using a Wald t-distribution with Kenward-Roger

approximation.

Emmeans (log-scale, scaled AUC)

contrast	estimate	SE	df lower.CL upper.CL t.ratio p.value

iMac - iMGL	1.59 0.34 2.02	0.145	3.04	4.687 0.0418

Results are averaged over the levels of: Line, Exp

Degrees-of-freedom method: kenward-roger

Confidence level used: 0.95

Back-transformed to AUC (iMac - iMGL): β = 3,671,297 AUC, 95% CI: [2,299,831, 5,384,776], P_adj = 0.042; fold change = 4.89 [2.51, 9.53]

Model assumptions

Warning: Non-normality of residuals detected (p = 0.016).

OK: Error variance appears to be homoscedastic (p = 0.798).

Singular fit (culture variance estimated at ~0): FALSE

\# Check for Multicollinearity

Low Correlation

Term VIF	VIF 95% CI adj. VIF Tolerance Tolerance 95% CI

Cell_Type 1.42 [1.12, 2.42]	1.19	0.71	[0.41, 0.89]

Line 1.61 [1.22, 2.68]	1.27	0.62	[0.37, 0.82]

Exp 1.15 [1.01, 2.70]	1.07	0.87	[0.37, 0.99]

Donor-dependence check (sensitivity analysis)

Cell type x line interaction: not estimable for this substrate

iMac - iMGL per line (model scale, 95% CI): not estimable - only 1 line(s) with both cell types (WTC-11)

AB-42

We fitted a linear mixed model (estimated using REML and nloptwrap optimizer) to predict

AB42_AUC_log with Cell_Type, Line and Exp (formula: AB42_AUC_log ~ Cell_Type + Line +

Exp). The model included Culture as random effect (formula: ~1 | Culture). The model's

total explanatory power is substantial (conditional R2 = 0.93) and the part related to

the fixed effects alone (marginal R2) is of 0.79. The model's intercept, corresponding to

Cell_Type = iMGL, Line = ID1 and Exp = 1.1, is at -1.36 (95% CI [-2.03, -0.69], t(6.87) =

-4.83, p = 0.002). Within this model:

- The effect of Cell Type [iMac] is statistically significant and positive (beta = 1.77,

95% CI [1.18, 2.36], t(7.17) = 7.00, p < .001; Std. beta = 1.77, 95% CI [1.18, 2.36])

- The effect of Line [KOLF] is statistically non-significant and positive (beta = 0.06,

95% CI [-0.70, 0.82], t(6.86) = 0.19, p = 0.855; Std. beta = 0.06, 95% CI [-0.70, 0.82])

- The effect of Line [WTC-11] is statistically non-significant and positive (beta = 0.63,

95% CI [-0.03, 1.29], t(7.10) = 2.26, p = 0.057; Std. beta = 0.63, 95% CI [-0.03, 1.29])

- The effect of Exp [3.1] is statistically non-significant and negative (beta = -0.45,

95% CI [-1.11, 0.21], t(7.19) = -1.60, p = 0.153; Std. beta = -0.45, 95% CI [-1.11,

0.21])

- The effect of Exp [4.1] is statistically non-significant and positive (beta = 0.58, 95%

CI [-0.16, 1.33], t(7.09) = 1.85, p = 0.106; Std. beta = 0.58, 95% CI [-0.16, 1.33])

Parameters were scaled by the mean and the SD of the response variable. 95% Confidence

Intervals (CIs) and p-values were computed using a Wald t-distribution with Kenward-Roger

approximation.

Emmeans (log-scale, scaled AUC)

contrast	estimate	SE	df lower.CL upper.CL t.ratio p.value

iMac - iMGL	1.77 0.253 7.17	1.18	2.36	7.003 0.0002

Results are averaged over the levels of: Line, Exp

Degrees-of-freedom method: kenward-roger

Confidence level used: 0.95

Back-transformed to AUC (iMac - iMGL): β = 77,751,223 AUC, 95% CI: [51,243,351, 114,088,289], P_adj = 3.8 × 10^-4; fold change = 7.9 [4.43, 14.3]

Model assumptions

OK: residuals appear as normally distributed (p = 0.396).

OK: Error variance appears to be homoscedastic (p = 0.954).

Singular fit (culture variance estimated at ~0): FALSE

\# Check for Multicollinearity

Low Correlation

Term VIF	VIF 95% CI adj. VIF Tolerance Tolerance 95% CI

Cell_Type 1.16 [1.02, 2.08]	1.08	0.86	[0.48, 0.98]

Line 1.25 [1.06, 2.01]	1.12	0.80	[0.50, 0.94]

Exp 1.25 [1.06, 2.01]	1.12	0.80	[0.50, 0.94]

Donor-dependence check (sensitivity analysis)

Cell type x line interaction (likelihood-ratio test, ML fits): chi2(2) = 1.24, P = 0.538

iMac - iMGL per line (model scale, 95% CI): ID1: 2.08 [0.92, 3.23]; KOLF: 1.87 [0.67, 3.07]; WTC-11: 1.60 [0.83, 2.38]

**Fig. 3E - Zymosan Phagocytosis**

Time Course

Normalised intensity was modelled with a generalised linear mixed model (glmmTMB; family: tweedie) with cell type, a natural spline of time (3 df) and their interaction, line and experiment as fixed effects; random intercepts for culture (grouping each culture's replicate wells) and a random intercept and time slope per well (repeated measures). The family was chosen from simulated-residual (DHARMa) diagnostics across compatible families; no transformation was applied. Cell type effects were tested by likelihood-ratio tests; iMac - iMGL differences at 4, 8, 12, 16, 20, 24 h are Holm-adjusted across timepoints.

Excluded wells: none

Family comparison

|family |converged | AIC| Uniformity_p| Dispersion_p| Outliers_p| ZeroInflation_p|

|tweedie |TRUE | 35981.85| 0| 0.056| 0.02| 0|

|hurdle_gamma |TRUE | 36698.61| 0| 0.000| 0.02| 1|

|gaussian_disp |FALSE | NA| 0| 0.000| 0.32| 0|

Likelihood-ratio tests

Time (main + interaction): chi2(6) = 866.49, P = 6.6 × 10^-184

Cell type x time interaction: chi2(3) = 154.12, P = 3.4 × 10^-33

Overall cell type effect (main + interaction): chi2(4) = 154.78, P = 1.9 × 10^-32

iMac - iMGL difference in normalised intensity by timepoint (Holm across timepoints)

| Elapsed| estimate| SE| asymp.LCL| asymp.UCL| p.value| p_adj|Significance |

| 4| -7631.603| 5957.349| -19307.793| 4044.587| 0.200| 0.200|ns |

| 8| 28922.051| 11018.375| 7326.432| 50517.670| 0.009| 0.020|* |

| 12| 59940.728| 14814.977| 30903.908| 88977.549| 0.000| 0.000|*** |

| 16| 64525.205| 16580.640| 32027.747| 97022.663| 0.000| 0.000|*** |

| 20| 63983.306| 19004.724| 26734.732| 101231.880| 0.001| 0.003|** |

| 24| 63394.707| 23380.300| 17570.160| 109219.254| 0.007| 0.020|* |

Fixed effects (model scale)

Estimate Std. Error z value Pr(>|z|)

(Intercept)	9.797	0.184 53.297 0.00e+00

Cell_TypeiMac	-0.819	0.147 -5.554 2.80e-08

ns(Elapsed, 3)1	0.589	0.092	6.406 1.50e-10

ns(Elapsed, 3)2	2.006	0.199 10.101 5.48e-24

ns(Elapsed, 3)3	0.498	0.112	4.466 7.96e-06

LineKOLF	-0.281	0.162 -1.739 8.20e-02

LineWTC-11	0.146	0.154	0.948 3.43e-01

Exp2.2	0.695	0.255	2.724 6.46e-03

Exp3.1	0.962	0.216	4.453 8.46e-06

Exp4.1	0.545	0.262	2.077 3.78e-02

Cell_TypeiMac:ns(Elapsed, 3)1	1.059	0.122	8.686 3.77e-18

Cell_TypeiMac:ns(Elapsed, 3)2	2.530	0.272	9.309 1.29e-20

Cell_TypeiMac:ns(Elapsed, 3)3	0.571	0.149	3.830 1.28e-04

Model diagnostics (DHARMa, chosen family)

Uniformity P = 1.4 × 10^-5; dispersion P = 0.056; outliers P = 0.020

AUC Values

Per-well area under the uptake curve (0-24 h, trapezoidal) was analysed untransformed, scaled, and modelled with a linear mixed model (fixed: cell type, line, experiment; random intercept: culture, grouping each culture's replicate wells), with Kenward-Roger degrees of freedom.

Excluded wells: none

Zymosan

We fitted a linear mixed model (estimated using REML and nloptwrap optimizer) to predict

Zymosan_AUC_identity with Cell_Type, Line and Exp (formula: Zymosan_AUC_identity ~

Cell_Type + Line + Exp). The model included Culture as random effect (formula: ~1 |

Culture). The model's total explanatory power is substantial (conditional R2 = 0.92) and

the part related to the fixed effects alone (marginal R2) is of 0.69. The model's

intercept, corresponding to Cell_Type = iMGL, Line = ID1 and Exp = 1.1, is at -1.35 (95%

CI [-2.10, -0.61], t(8.92) = -4.11, p = 0.003). Within this model:

- The effect of Cell Type [iMac] is statistically significant and positive (beta = 0.70,

95% CI [0.08, 1.31], t(8.99) = 2.55, p = 0.031; Std. beta = 0.70, 95% CI [0.08, 1.31])

- The effect of Line [KOLF] is statistically non-significant and negative (beta = -0.25,

95% CI [-1.09, 0.58], t(8.91) = -0.69, p = 0.509; Std. beta = -0.25, 95% CI [-1.09,

0.58])

- The effect of Line [WTC-11] is statistically non-significant and positive (beta = 0.27,

95% CI [-0.53, 1.06], t(9.06) = 0.76, p = 0.466; Std. beta = 0.27, 95% CI [-0.53, 1.06])

- The effect of Exp [2.2] is statistically significant and positive (beta = 1.16, 95% CI

[0.06, 2.27], t(9.07) = 2.38, p = 0.041; Std. beta = 1.16, 95% CI [0.06, 2.27])

- The effect of Exp [3.1] is statistically significant and positive (beta = 1.69, 95% CI

[0.94, 2.43], t(8.98) = 5.13, p < .001; Std. beta = 1.69, 95% CI [0.94, 2.43])

- The effect of Exp [4.1] is statistically significant and positive (beta = 1.15, 95% CI

[0.24, 2.06], t(9.08) = 2.84, p = 0.019; Std. beta = 1.15, 95% CI [0.24, 2.06])

Parameters were scaled by the mean and the SD of the response variable. 95% Confidence

Intervals (CIs) and p-values were computed using a Wald t-distribution with Kenward-Roger

approximation.

Emmeans (identity-scale, scaled AUC)

contrast	estimate	SE	df lower.CL upper.CL t.ratio p.value

iMac - iMGL	0.696 0.273 8.99	0.0779	1.31	2.548 0.0313

Results are averaged over the levels of: Line, Exp

Degrees-of-freedom method: kenward-roger

Confidence level used: 0.95

Back-transformed to AUC (iMac - iMGL): β = 1,251,676 AUC, 95% CI: [278,663, 2,217,516], P = 0.031; fold change = 1.57 [1.1, 2.48]

Model assumptions

OK: residuals appear as normally distributed (p = 0.425).

OK: Error variance appears to be homoscedastic (p = 0.820).

Singular fit (culture variance estimated at ~0): FALSE

\# Check for Multicollinearity

Low Correlation

Term VIF	VIF 95% CI adj. VIF Tolerance Tolerance 95% CI

Cell_Type 1.02 [1.00, 305.18]	1.01	0.98	[0.00, 1.00]

Line 1.48 [1.20, 2.14]	1.22	0.67	[0.47, 0.83]

Exp 1.50 [1.22, 2.17]	1.23	0.66	[0.46, 0.82]

Donor-dependence check (sensitivity analysis)

Cell type x line interaction (likelihood-ratio test, ML fits): chi2(2) = 0.06, P = 0.970

iMac - iMGL per line (model scale, 95% CI): ID1: 0.62 [-0.54, 1.78]; KOLF: 0.75 [-0.48, 1.98]; WTC-11: 0.72 [-0.24, 1.68]

**Figure 3G and H - Lysosomal Mass and Acidification**

gMFI values were analysed untransformed unless residual diagnostics indicated that model assumptions were not met (transformation used: LysoSensor gMFI = log; LysoTracker gMFI = log), scaled, and modelled per marker with a linear mixed model (fixed: Cell_Type, Donor, Experiment; random intercept: culture, grouping each culture's replicate wells), with Kenward-Roger degrees of freedom. P values within each model are unadjusted; P_adj is Holm-adjusted across the 2 markers.

Excluded rows: none

Design (wells per cell type, donor and experiment)

|Cell_Type |Donor |Experiment | n|

|iMGL |ID1 |1.1 | 3|

|iMGL |ID1 |2.1 | 3|

|iMGL |ID1 |3.1 | 3|

|iMGL |KOLF |1.1 | 3|

|iMGL |KOLF |2.1 | 3|

|iMGL |KOLF |3.1 | 3|

|iMGL |WTC-11 |1.1 | 3|

|iMGL |WTC-11 |2.1 | 3|

|iMGL |WTC-11 |2.2 | 3|

|iMGL |WTC-11 |4.1 | 2|

|iMac |ID1 |1.1 | 3|

|iMac |ID1 |2.1 | 3|

|iMac |ID1 |3.1 | 3|

|iMac |ID1 |4.1 | 2|

|iMac |KOLF |1.1 | 3|

|iMac |KOLF |2.1 | 3|

|iMac |KOLF |2.2 | 3|

|iMac |KOLF |3.1 | 3|

|iMac |WTC-11 |1.1 | 1|

|iMac |WTC-11 |2.2 | 3|

|iMac |WTC-11 |3.1 | 3|

|iMac |WTC-11 |4.1 | 2|

LysoSensor gMFI

We fitted a linear mixed model (estimated using REML and nloptwrap optimizer) to predict

LysoSensor_log with Cell_Type, Donor and Experiment (formula: LysoSensor_log ~ Cell_Type

+ Donor + Experiment). The model included Culture as random effect (formula: ~1 |

Culture). The model's total explanatory power is substantial (conditional R2 = 0.74) and

the part related to the fixed effects alone (marginal R2) is of 0.71. The model's

intercept, corresponding to Cell_Type = iMGL, Donor = ID1 and Experiment = 1.1, is at

1.47 (95% CI [1.03, 1.90], t(12.70) = 7.27, p < .001). Within this model:

- The effect of Cell Type [iMac] is statistically non-significant and positive (beta =

3.37e-04, 95% CI [-0.35, 0.35], t(13.74) = 2.09e-03, p = 0.998; Std. beta = 3.37e-04, 95%

CI [-0.35, 0.35])

- The effect of Donor [KOLF] is statistically non-significant and negative (beta = -0.06,

95% CI [-0.47, 0.36], t(12.70) = -0.29, p = 0.776; Std. beta = -0.06, 95% CI [-0.47,

0.36])

- The effect of Donor [WTC-11] is statistically non-significant and negative (beta =

-0.23, 95% CI [-0.68, 0.22], t(14.59) = -1.09, p = 0.293; Std. beta = -0.23, 95% CI

[-0.68, 0.22])

- The effect of Experiment [2.1] is statistically significant and negative (beta = -2.19,

95% CI [-2.67, -1.72], t(12.86) = -10.08, p < .001; Std. beta = -2.19, 95% CI [-2.67,

-1.72])

- The effect of Experiment [2.2] is statistically significant and negative (beta = -1.63,

95% CI [-2.21, -1.05], t(13.57) = -6.04, p < .001; Std. beta = -1.63, 95% CI [-2.21,

-1.05])

- The effect of Experiment [3.1] is statistically significant and negative (beta = -1.72,

95% CI [-2.19, -1.24], t(13.04) = -7.84, p < .001; Std. beta = -1.72, 95% CI [-2.19,

-1.24])

- The effect of Experiment [4.1] is statistically significant and negative (beta = -1.73,

95% CI [-2.34, -1.11], t(19.50) = -5.83, p < .001; Std. beta = -1.73, 95% CI [-2.34,

-1.11])

Parameters were scaled by the mean and the SD of the response variable. 95% Confidence

Intervals (CIs) and p-values were computed using a Wald t-distribution with Kenward-Roger

approximation.

Emmeans (log-scale, scaled gMFI)

contrast	estimate	SE	df lower.CL upper.CL t.ratio p.value

iMac - iMGL 0.000337 0.161 13.7	-0.347	0.347	0.002 0.9984

Results are averaged over the levels of: Donor, Experiment

Degrees-of-freedom method: kenward-roger

Confidence level used: 0.95

Back-transformed to gMFI (iMac - iMGL): β = 0 gMFI, 95% CI: [-156, 152], P_adj = 0.998; fold change = 1 [0.741, 1.36]

Model assumptions

Warning: Non-normality of residuals detected (p < .001).

OK: Error variance appears to be homoscedastic (p = 0.396).

Singular fit (culture variance estimated at ~0): FALSE

\# Check for Multicollinearity

Low Correlation

Term VIF	VIF 95% CI adj. VIF Tolerance Tolerance 95% CI

Cell_Type 1.09 [1.01, 2.44]	1.04	0.92	[0.41, 0.99]

Donor 1.30 [1.10, 1.92]	1.14	0.77	[0.52, 0.91]

Experiment 1.36 [1.13, 1.97]	1.16	0.74	[0.51, 0.88]

Donor-dependence check (sensitivity analysis)

Cell type x donor interaction (likelihood-ratio test, ML fits): chi2(2) = 0.41, P = 0.813

iMac - iMGL per donor (model scale, 95% CI): ID1: 0.08 [-0.52, 0.69]; KOLF: -0.12 [-0.72, 0.48]; WTC-11: 0.03 [-0.57, 0.64]

LysoTracker gMFI

We fitted a linear mixed model (estimated using REML and nloptwrap optimizer) to predict

LysoTracker_log with Cell_Type, Donor and Experiment (formula: LysoTracker_log ~

Cell_Type + Donor + Experiment). The model included Culture as random effect (formula: ~1

| Culture). The model's total explanatory power is substantial (conditional R2 = 0.97)

and the part related to the fixed effects alone (marginal R2) is of 0.78. The model's

intercept, corresponding to Cell_Type = iMGL, Donor = ID1 and Experiment = 1.1, is at

0.39 (95% CI [-0.18, 0.97], t(13.87) = 1.46, p = 0.166). Within this model:

- The effect of Cell Type [iMac] is statistically significant and positive (beta = 1.22,

95% CI [0.78, 1.67], t(14.04) = 5.91, p < .001; Std. beta = 1.22, 95% CI [0.78, 1.67])

- The effect of Donor [KOLF] is statistically non-significant and negative (beta = -0.20,

95% CI [-0.75, 0.35], t(13.86) = -0.79, p = 0.445; Std. beta = -0.20, 95% CI [-0.75,

0.35])

- The effect of Donor [WTC-11] is statistically non-significant and positive (beta =

0.55, 95% CI [-0.02, 1.11], t(14.14) = 2.08, p = 0.057; Std. beta = 0.55, 95% CI [-0.02,

1.11])

- The effect of Experiment [2.1] is statistically significant and negative (beta = -1.65,

95% CI [-2.26, -1.03], t(13.93) = -5.76, p < .001; Std. beta = -1.65, 95% CI [-2.26,

-1.03])

- The effect of Experiment [2.2] is statistically significant and negative (beta = -1.25,

95% CI [-1.99, -0.51], t(14.07) = -3.61, p = 0.003; Std. beta = -1.25, 95% CI [-1.99,

-0.51])

- The effect of Experiment [3.1] is statistically significant and negative (beta = -1.57,

95% CI [-2.19, -0.96], t(13.97) = -5.51, p < .001; Std. beta = -1.57, 95% CI [-2.19,

-0.96])

- The effect of Experiment [4.1] is statistically significant and negative (beta = -1.72,

95% CI [-2.46, -0.97], t(14.42) = -4.94, p < .001; Std. beta = -1.72, 95% CI [-2.46,

-0.97])

Parameters were scaled by the mean and the SD of the response variable. 95% Confidence

Intervals (CIs) and p-values were computed using a Wald t-distribution with Kenward-Roger

approximation.

Emmeans (log-scale, scaled gMFI)

contrast	estimate	SE df lower.CL upper.CL t.ratio p.value

iMac - iMGL	1.22 0.207 14	0.78	1.67	5.911 <.0001

Results are averaged over the levels of: Donor, Experiment

Degrees-of-freedom method: kenward-roger

Confidence level used: 0.95

Back-transformed to gMFI (iMac - iMGL): β = 587 gMFI, 95% CI: [369, 871], P_adj = 7.5 × 10^-5; fold change = 3.95 [2.51, 6.24]

Model assumptions

OK: residuals appear as normally distributed (p = 0.578).

OK: Error variance appears to be homoscedastic (p = 0.855).

Singular fit (culture variance estimated at ~0): FALSE

\# Check for Multicollinearity

Low Correlation

Term VIF	VIF 95% CI adj. VIF Tolerance Tolerance 95% CI

Cell_Type 1.07 [1.00, 3.38]	1.03	0.94	[0.30, 1.00]

Donor 1.28 [1.09, 1.90]	1.13	0.78	[0.53, 0.92]

Experiment 1.33 [1.12, 1.94]	1.15	0.75	[0.51, 0.90]

Donor-dependence check (sensitivity analysis)

Cell type x donor interaction (likelihood-ratio test, ML fits): chi2(2) = 5.26, P = 0.072

iMac - iMGL per donor (model scale, 95% CI): ID1: 1.72 [1.01, 2.42]; KOLF: 1.14 [0.44, 1.85]; WTC-11: 0.86 [0.20, 1.52]

**Fig. 4B - TREM2 CTF/FL Western Blots**

WT iMGL and iMac membrane fractions were run side by side on the same blots; only such blots were analysed. Band intensity was analysed untransformed unless residual diagnostics indicated that model assumptions were not met (transformation used: TREM2 FL = identity; TREM2 CTF = identity; the Na/K-ATPase covariate received the same transformation), scaled, and modelled with cell type, Na/K-ATPase intensity on the same lane (covariate) and blot (fixed effect with fewer than 5 blots; random intercept otherwise). The iMac - iMGL difference is given at the mean Na/K-ATPase intensity, back-transformed to band-intensity units, with 95% CI and fold difference; P_adj is Holm-adjusted across TREM2-FL and TREM2-CTF. Treatment condition of the lanes analysed (from the sample key): iMGL = IL-4; iMac = IL-4.

Design (lanes per band, blot and cell type)

|Antibody |Blot | iMGL| iMac|

|TREM2 CTF |20240923_1 | 1| 1|

|TREM2 CTF |20250910_1 | 3| 2|

|TREM2 FL |20240923_1 | 1| 1|

|TREM2 FL |20250910_1 | 3| 2|

TREM2 FL

Model: lm, blot as fixed effect; transformation: identity

Parameter	| Coefficient |	SE |	95% CI | t(3) |	p

----------------------------------------------------------------------

(Intercept)	|	-0.64 | 0.27 | [-1.51, 0.22] | -2.38 | 0.098

Cell Type [iMac] |	1.86 | 0.26 | [ 1.04, 2.68] | 7.20 | 0.006

NaK	|	0.10 | 0.16 | [-0.42, 0.63] | 0.63 | 0.571

Blot [20250910_1] |	-0.21 | 0.32 | [-1.23, 0.80] | -0.67 | 0.550

iMac - iMGL: β = 11,513 (band intensity), 95% CI: [8,308, 14,652]; 8.00-fold (95% CI: -55.20–65.70), P_adj = 0.011; unadjusted P = 0.006, residual df = 3.00

Model assumptions (few observations - interpret with caution)

OK: residuals appear as normally distributed (p = 0.411).

OK: Error variance appears to be homoscedastic (p = 0.184).

TREM2 CTF

Model: lm, blot as fixed effect; transformation: identity

Parameter	| Coefficient |	SE |	95% CI | t(3) |	p

----------------------------------------------------------------------

(Intercept)	|	-0.38 | 0.31 | [-1.38, 0.62] | -1.21 | 0.313

Cell Type [iMac] |	1.89 | 0.30 | [ 0.94, 2.84] | 6.31 | 0.008

NaK	|	0.37 | 0.19 | [-0.23, 0.98] | 1.97 | 0.143

Blot [20250910_1] |	-0.60 | 0.37 | [-1.78, 0.58] | -1.62 | 0.203

iMac - iMGL: β = 10,857 (band intensity), 95% CI: [7,456, 14,253]; 4.05-fold (95% CI: 2.38–10.70), P_adj = 0.011; unadjusted P = 0.008, residual df = 3.00

Model assumptions (few observations - interpret with caution)

OK: residuals appear as normally distributed (p = 0.409).

OK: Error variance appears to be homoscedastic (p = 0.663).

**Fig. 5C p-SYK and p-PLCG2 Western Blots**

For each phospho-protein, band intensity (WT only; IgG control vs TREM2 crosslinking, 5 min) was log-transformed so that the cell type x treatment interaction estimates the ratio of fold inductions (residual diagnostics checked; transformation used: P-SYK = log; P-PLCg2 = log; loading-control covariate: same transformation as the band), and modelled with a linear mixed model (fixed: cell type, treatment, their interaction, and total-protein (SYK or PLCg2) intensity on the same lane; random intercepts: culture and IgG/TREM2 lane pair), with Kenward-Roger degrees of freedom. iMGLs and iMacs were run on separate blots, so cell types are compared by fold induction (TREM2 vs IgG within each lane pair), not by absolute band intensity. The cell type x treatment interaction is reported as the ratio of fold inductions (iMac / iMGL); P_adj is Holm-adjusted across 2 phospho-proteins.

Design (lanes per protein, cell type and treatment)

|Antibody |Cell_Type | IgG 5 min| TREM2 5 min|

|P-PLCg2 |iMGL | 6| 6|

|P-PLCg2 |iMac | 5| 5|

|P-SYK |iMGL | 9| 9|

|P-SYK |iMac | 9| 9|

P-SYK

We fitted a linear mixed model (estimated using REML and nloptwrap optimizer) to predict

P.SYK_log with Cell_Type, Treatment and LC (formula: P.SYK_log ~ Cell_Type * Treatment +

LC). The model included Culture as random effects (formula: list(~1 | Culture, ~1 |

Pair)). The model's explanatory power related to the fixed effects alone (marginal R2) is

0.69. The model's intercept, corresponding to Cell_Type = iMGL, Treatment = IgG 5 min and

LC = 0, is at 0.33 (95% CI [-4.70, 5.35], t(24.87) = 0.13, p = 0.895). Within this model:

- The effect of Cell Type [iMac] is statistically non-significant and negative (beta =

-0.67, 95% CI [-1.39, 0.04], t(18.61) = -1.98, p = 0.063; Std. beta = -0.78, 95% CI

[-1.61, 0.05])

- The effect of Treatment [TREM2 5 min] is statistically significant and positive (beta =

0.85, 95% CI [0.44, 1.27], t(15.95) = 4.36, p < .001; Std. beta = 0.99, 95% CI [0.51,

1.47])

- The effect of LC is statistically significant and positive (beta = 0.83, 95% CI [0.25,

1.42], t(24.50) = 2.93, p = 0.007; Std. beta = 0.37, 95% CI [0.11, 0.63])

- The effect of Cell Type [iMac] × Treatment [TREM2 5 min] is statistically

non-significant and positive (beta = 0.53, 95% CI [-0.02, 1.09], t(15.07) = 2.07, p =

0.056; Std. beta = 0.62, 95% CI [-0.02, 1.26])

Standardized parameters were obtained by fitting the model on a standardized version of

the dataset. 95% Confidence Intervals (CIs) and p-values were computed using a Wald

t-distribution with Kenward-Roger approximation.

Fold induction (TREM2 vs IgG): iMGL 2.35 [1.59, 3.46], P_adj = 9.9 × 10^-4; iMac 4.00 [2.69, 5.90], P_adj = 1.2 × 10^-5

Cell type x treatment interaction (ratio of fold inductions, iMac / iMGL): 1.71 [1.02, 2.83], t(15.07), P = 0.056, P_adj = 0.056

Main text: fold induction iMGL 2.35 (95% CI: 1.59–3.46), iMac 4.00 (95% CI: 2.69–5.90); iMac/iMGL ratio = 1.71, 95% CI: 1.02–2.83, P_adj = 0.056

Model assumptions

OK: residuals appear as normally distributed (p = 0.286).

OK: Error variance appears to be homoscedastic (p = 0.878).

Singular fit (a random-effect variance estimated at ~0): TRUE

P-PLCg2

We fitted a linear mixed model (estimated using REML and nloptwrap optimizer) to predict

P.PLCg2_log with Cell_Type, Treatment and LC (formula: P.PLCg2_log ~ Cell_Type *

Treatment + LC). The model included Culture as random effects (formula: list(~1 |

Culture, ~1 | Pair)). The model's total explanatory power is substantial (conditional R2

= 0.85) and the part related to the fixed effects alone (marginal R2) is of 0.71. The

model's intercept, corresponding to Cell_Type = iMGL, Treatment = IgG 5 min and LC = 0,

is at 0.47 (95% CI [-7.93, 8.86], t(16.93) = 0.12, p = 0.908). Within this model:

- The effect of Cell Type [iMac] is statistically significant and negative (beta = -2.39,

95% CI [-3.53, -1.25], t(13.83) = -4.50, p < .001; Std. beta = -1.57, 95% CI [-2.31,

-0.82])

- The effect of Treatment [TREM2 5 min] is statistically non-significant and positive

(beta = 0.80, 95% CI [-0.03, 1.63], t(9.05) = 2.19, p = 0.056; Std. beta = 0.53, 95% CI

[-0.02, 1.07])

- The effect of LC is statistically non-significant and positive (beta = 0.78, 95% CI

[-0.26, 1.83], t(16.94) = 1.58, p = 0.133; Std. beta = 0.22, 95% CI [-0.08, 0.52])

- The effect of Cell Type [iMac] × Treatment [TREM2 5 min] is statistically significant

and positive (beta = 2.10, 95% CI [0.86, 3.33], t(9.16) = 3.83, p = 0.004; Std. beta =

1.37, 95% CI [0.56, 2.18])

Standardized parameters were obtained by fitting the model on a standardized version of

the dataset. 95% Confidence Intervals (CIs) and p-values were computed using a Wald

t-distribution with Kenward-Roger approximation.

Fold induction (TREM2 vs IgG): iMGL 2.23 [1.08, 4.58], P_adj = 0.056; iMac 18.17 [8.61, 39.32], P_adj = 1.5 × 10^-4

Cell type x treatment interaction (ratio of fold inductions, iMac / iMGL): 8.15 [2.83, 23.86], t(9.16), P = 0.004, P_adj = 0.008

Main text: fold induction iMGL 2.23 (95% CI: 1.08–4.58), iMac 18.20 (95% CI: 8.61–39.32); iMac/iMGL ratio = 8.15, 95% CI: 2.83–23.86, P_adj = 0.008

Model assumptions

OK: residuals appear as normally distributed (p = 0.499).

OK: Error variance appears to be homoscedastic (p = 0.989).

Singular fit (a random-effect variance estimated at ~0): FALSE

**Fig. 5B,C - MEF2C KD in iMGLs and iMacs**

MEF2C band intensity was analysed after log transformation (untransformed data did not meet model assumptions), with actin on the same lane as a covariate (same transformation); both were scaled and modelled with a linear mixed model (fixed: genotype (SCR vs MEF2C-KD), cell type, their interaction and actin; random intercepts: culture (donor x cell type)), with Kenward-Roger degrees of freedom. Knockdown (MEF2C-KD vs SCR) was tested within each cell type, Holm-adjusted across the two cell types, and is reported as % reduction in MEF2C at the mean actin level (back-transformed). The genotype x cell type interaction tests whether knockdown differs between iMacs and iMGLs.

Design (lanes per cell type, donor and genotype)

|Cell_type |Donor | SCR| MEF2C-KD|

|iMGL |ADRC5 | 2| 2|

|iMGL |F12455 | 1| 1|

|iMGL |KOLF2.1J | 6| 6|

|iMGL |WTC11 | 2| 2|

|iMac |ADRC5 | 1| 1|

|iMac |KOLF2.1J | 1| 1|

|iMac |WTC11 | 1| 1|

|iMac |x051118 | 1| 1|

MEF2C Band Intensity

MEF2C band intensity was analysed after log transformation (untransformed data did not meet model assumptions), with actin on the same lane as a covariate (same transformation); both were scaled and modelled with a linear mixed model (fixed: genotype (SCR vs MEF2C-KD), cell type, their interaction and actin; random intercepts: culture (donor x cell type)), with Kenward-Roger degrees of freedom. Knockdown (MEF2C-KD vs SCR) was tested within each cell type, Holm-adjusted across the two cell types, and is reported as % reduction in MEF2C at the mean actin level (back-transformed). The genotype x cell type interaction tests whether knockdown differs between iMacs and iMGLs.

Design (lanes per cell type, donor and genotype)

|Cell_type |Donor | SCR| MEF2C-KD|

|iMGL |ADRC5 | 2| 2|

|iMGL |F12455 | 1| 1|

|iMGL |KOLF2.1J | 6| 6|

|iMGL |WTC11 | 2| 2|

|iMac |ADRC5 | 1| 1|

|iMac |KOLF2.1J | 1| 1|

|iMac |WTC11 | 1| 1|

|iMac |x051118 | 1| 1|

MEF2C KD Efficiency

We fitted a linear mixed model (estimated using REML and nloptwrap optimizer) to predict

MEF2C_log with Genotype, Cell_type and Actin_c (formula: MEF2C_log ~ Genotype * Cell_type

+ Actin_c). The model included Culture as random effect (formula: ~1 | Culture). The

model's total explanatory power is substantial (conditional R2 = 0.84) and the part

related to the fixed effects alone (marginal R2) is of 0.78. The model's intercept,

corresponding to Genotype = SCR, Cell_type = iMGL and Actin_c = 0, is at 0.57 (95% CI

[0.12, 1.02], t(7.15) = 2.96, p = 0.021). Within this model:

- The effect of Genotype [MEF2C-KD] is statistically significant and negative (beta =

-0.63, 95% CI [-1.00, -0.26], t(19.54) = -3.52, p = 0.002; Std. beta = -0.63, 95% CI

[-1.00, -0.26])

- The effect of Cell type [iMac] is statistically non-significant and negative (beta =

-0.29, 95% CI [-0.98, 0.39], t(13.20) = -0.93, p = 0.372; Std. beta = -0.29, 95% CI

[-0.98, 0.39])

- The effect of Actin c is statistically significant and positive (beta = 0.57, 95% CI

[0.38, 0.76], t(22.84) = 6.24, p < .001; Std. beta = 0.57, 95% CI [0.38, 0.76])

- The effect of Genotype [MEF2C-KD] × Cell type [iMac] is statistically significant and

negative (beta = -0.73, 95% CI [-1.44, -0.03], t(19.35) = -2.18, p = 0.042; Std. beta =

-0.73, 95% CI [-1.44, -0.03])

Standardized parameters were obtained by fitting the model on a standardized version of

the dataset. 95% Confidence Intervals (CIs) and p-values were computed using a Wald

t-distribution with Kenward-Roger approximation.

Knockdown within cell type (model scale; KD - SCR)

Cell_type = iMGL:

contrast	estimate	SE	df lower.CL upper.CL t.ratio p.value

(MEF2C-KD) - SCR	-0.63 0.179 19.5	-1.00	-0.256 -3.517 0.0022

Cell_type = iMac:

contrast	estimate	SE	df lower.CL upper.CL t.ratio p.value

(MEF2C-KD) - SCR	-1.36 0.286 19.3	-1.96	-0.767 -4.777 0.0001

Degrees-of-freedom method: kenward-roger

Confidence level used: 0.95

iMac: 72.0% reduction in MEF2C (95% CI: 53.1–83.5%), P_adj = 2.5 × 10^-4 (unadjusted P = 1.3 × 10^-4)

iMGL: 44.4% reduction in MEF2C (95% CI: 23.0–59.6%), P_adj = 0.002 (unadjusted P = 0.002)

Genotype x cell type interaction: estimate = -0.73 (model scale), 95% CI [-1.44, -0.03], t(19.3), P = 0.042

Model assumptions

OK: residuals appear as normally distributed (p = 0.120).

OK: Error variance appears to be homoscedastic (p = 0.997).

Singular fit (a random-effect variance estimated at ~0): FALSE

\# Check for Multicollinearity

Low Correlation

Term VIF	VIF 95% CI adj. VIF Tolerance Tolerance 95% CI

Genotype 1.47 [1.15, 2.45]	1.21	0.68	[0.41, 0.87]

Cell_type 1.57 [1.20, 2.59]	1.25	0.64	[0.39, 0.83]

Actin_c 1.14 [1.01, 2.69]	1.07	0.88	[0.37, 0.99]

Genotype:Cell_type 1.85 [1.36, 3.02]	1.36	0.54	[0.33, 0.74]

MEF2C KD Efficiency by Donor

To test whether knockdown efficiency differed between donors, the model above was extended with donor and genotype x donor as fixed effects (all lanes included; same transformation, covariate and random intercept). Donor differences in knockdown were tested with a Kenward-Roger F-test comparing this model with the same model without the genotype x donor terms. Because each iMac donor contributed a single SCR/KD pair, donor efficiency was assumed to be the same in both cell types in the pooled test; an iMGL-only test was also performed (ordinary F-test, as culture and donor coincide within iMGL), and an iMac-only test is not estimable.

Genotype x donor (pooled): F(4, 15.0) = 1.20, P = 0.353
 Genotype x donor (iMGL only): F(3, 13) = 2.30, P = 0.126

Knockdown per donor (% reduction, 95% CI, Holm-adjusted across 8 tests):
 iMac: ADRC5 83.4% (64.8–92.2%), P_adj = 0.003; KOLF2.1J 68.2% (35.4–84.5%), P_adj = 0.036; WTC11 72.3% (40.3–87.3%), P_adj = 0.030; x051118 57.6% (−21.7–85.3%), P_adj = 0.256
 iMGL: ADRC5 67.9% (38.3–83.0%), P_adj = 0.027; F12455 10.2% (−155.8–68.6%), P_adj = 0.844; KOLF2.1J 38.3% (6.1–59.7%), P_adj = 0.160; WTC11 46.3% (−2.4–71.6%), P_adj = 0.241
